# Beyond functional connectivity: deep learning applied to resting-state fMRI time series in the prediction of 58 human traits in the HCP

**DOI:** 10.1101/2024.03.07.583858

**Authors:** Bruno Hebling Vieira, Mikkel Schöttner, Vince D Calhoun, Carlos Ernesto Garrido Salmon

## Abstract

Machine learning has made several inroads into the study of brain-behavior relations based on in vivo imaging. While the advent of deep neural networks was expected to further improve predictions, the current literature based on resting-state functional connectivity presents mixed results. We hypothesize that the representation of the data, *i.e.* in the form of functional connectivity, could restrict an advantage of deep learning techniques, namely that of learning complex representations directly from the data. Thus, we investigated if bypassing this feature extraction resulted in improved performance in the prediction of 58 widely studied behavioral traits from a large sample of Human Connectome Project subjects, using deep learning techniques. For this task, we adapted the InceptionTime architecture, which jointly predicts traits directly from regional time series through representation learning, and compared results with a strong kernel-based baseline. Results revealed that both models achieve comparable performance in most traits. Eleven significant differences in mean squared error were detected, however, with seven favoring the neural network approach, and this number increased when accounting for covariates. We additionally show that contrary to the expectation, the neural network approach was more robust to reductions in the training set size. On the other hand, it was more sensitive to reductions in the length of the time series at test time. Our results present a more nuanced view of the potential of deep learning for the prediction of behavior from neuroimaging, which allows learning features directly from the data.

## 1 Introduction

Neuroimaging-based prediction of behavior using machine learning is of great interest in the literature [1–14]. It is a natural extension of the study of group-level brain-behavior relations, which has been a recurring topic in contemporary cognitive neuroscience [15]. The advent of machine learning has enabled the individualized prediction of complex multivariate relations between brain and behavior. As shown in Sui et al. [16] and Vieira et al. [17], the number of works being published in this area of research is currently undergoing accelerated expansion. Most of the literature employs traditional approaches to predict behavior, such as kernel ridge regression (KRR) and connectome predictive modeling. Table 1 demonstrates the recent emergence of studies that simultaneously investigate the prediction of multiple behavioral traits using machine learning from magnetic resonance imaging (MRI)-extracted data.

**Table 1:** Non-exhaustive list of studies that investigate machine learning based prediction of diverse traits from MRI. The number of traits is reported, with the corresponding number of subjects. ^†^: In all eight cases, the 58 traits in the HCP refer to the traits presented in Table S1. Other studies employ different sets of traits, with possible overlaps. CPM: connectome predictive modeling. SVR: support vector regression. PLSR: partial least squares regression. PCR: principal component regression. GSP: Brain Genomics Superstruct Project. UKBB: UK Biobank. ABCD: Adolescent Brain Cognitive Development study. LRR: linear ridge regression. FC: functional connectivity. SC: structural connectivity. TBSS: tract-based spatial statistics.

| Studies | Year | N | Traits | Inputs | Model |
| --- | --- | --- | --- | --- | --- |
| **HCP fMRI** |  |  |  |  |  |
| Kong et al. [4] | 2019 | 577 | 58<sup>†</sup> | fMRI network<br>spatial topography | KRR |
| Kashyap et al. [5] | 2019 | 803 | 58<sup>†</sup> | RSFC | Elastic-Net |
| Li et al. [6] | 2019 | 953 | 58<sup>†</sup> | RSFC | KRR |
| He et al. [7] | 2020 | 953 | 58<sup>†</sup> | RSFC | KRR, BrainNetCNN,<br>GCNN, FNN |
| Lin et al. [8] | 2020 | 144 | 29 | RSFC | CPM |
| Wei et al. [9] | 2020 | 1003 | 13 | RSFC | CPM, SVR, LASSO,<br>Ridge |
| Wu et al. [10] | 2020 | 922 | 98 | Task fMRI<br>statistical maps | PLSR |
| Kong et al. [11] | 2021 | 752 | 58<sup>†</sup> | RSFC | KRR |
| Li et al. [12] | 2022 | 948 | 58<sup>†</sup> | RSFC | KRR |
| Ooi et al. [13] | 2022 | 753 | 58<sup>†</sup> | RSFC, task FC<br>(plus morphometry,<br>SC, and TBSS) | KRR, LRR, Elastic-Net<br>regression |
| Kong et al. [14] | 2023 | 746 | 58<sup>†</sup> | RSFC | KRR, LRR |
| **Other fMRI** |  |  |  |  |  |
| Li et al. [6]<br>(GSP) | 2019 | 862 | 23 | RSFC | KRR |
| He et al. [7]<br>(UKBB) | 2020 | 8868 | 4 | RSFC | KRR, BrainNetCNN,<br>GCNN, FNN |
| Li et al. [12]<br>(ABCD) | 2022 | 5351 | 36 | RSFC | KRR |
| Chen et al. [18]<br>(ABCD) | 2022 | 1858 | 36 | RSFC, task FC | KRR |
| Ooi et al. [13]<br>(ABCD) | 2022 | 1823 | 36 | RSFC, task FC<br>(plus morphometry,<br>SC, and TBSS) | KRR, LRR, Elastic-Net<br>regression |
| Kong et al. [14]<br>(ABCD) | 2023 | 1476 | 36 | RSFC | KRR, LRR |

Potential limitations of traditional methods, such as the need of engineering features relating brain and behavior, have motivated the use of deep neural networks to predict behavior from neuroimaging [7, 19, 20]. However, it has been demonstrated that newer deep-learning-based approaches using resting-state functional connectivity (RSFC) as input do not outperform classical KRR on the prediction of behavior and demographics [7]. On the other hand, deep-learning was shown to be able to learn abstract features from anatomical data, showing great potential when compared with traditional methods [21]. Deep neural networks were also successfully used in the prediction of intelligence from functional MRI (fMRI) time series [19, 20]. None of the studies presented in Table 1 used fMRI time series directly for prediction, instead relying on region of interest (ROI)-level features. It is not clear how deep-learning compares to traditional methods when trained on lower level, structured, fMRI data for the prediction of diverse behavioral traits. Additionally, no evidence has been found for different scaling of performance as a function of available training data between RSFC-based deep learning and traditional models in the prediction of several traits in the UK Biobank [7, 22]. Or, in other words, the performance of the deep learning models did not differ from that observed in the traditional models across different training set sizes. These previous findings, where no differences in performance were found, point towards a possible bottleneck in the data itself, *i.e.* lack of non-linearities in the relation between RSFC and behavior that could be exploited by deep learning. Therefore, the scaling of fMRI time series-based deep learning on both the number of subjects during training and the number of time points available at test time has not yet been established. It is possible that different scaling exists between RSFC-based traditional methods and time series-based deep learning.

To address these questions, we performed a study comparing performance between a traditional RSFC-based model and a fMRI time series-based deep learning model across the same 58 behavioral variables in the Human Connectome Project (HCP), previously studied in multiple studies [4–7, 11–14]. For the traditional method, we implemented the KRR based on the RSFC matrices in the HCP, as in He et al. [7]. This is a well-established method that sees widespread usage in the literature, as can be attested according to Table 1. For the deep-learning method based on cortical time series, we adopted the InceptionTime network, a temporal convolutional neural network (TCNN) specially tailored for supervised-learning with time series [23]. A TCNN offers several advantages over other time series-based deep-learning approaches, such as recurrent neural networks (RNNs), including the ability to process longer time series and the ability to parallelize computation, which is important for the large number of subjects in the HCP dataset. We demonstrate significant differences in performance between the two methods, with the deep-learning approach attaining lower error in 7 out of 58 traits, and higher error in 4 out of 58 traits. We also show that the models scale differently. While KRR is more sensitive to the number of subjects in the training set, on the other hand the TCNN performance is largely unaffected when training on smaller subsamples, although the fully trained TCNN does not generalize as well to shorter time series.

## 2 Methods

### 2.1 Resting-state fMRI acquisition

HCP resting-state fMRI (RS-fMRI) data acquisition was performed on a customized Siemens 3 Tesla scanner. Participants were scanned in two days with two runs on each day. Runs alternated between right-to-left and left-to-right phase encoding, to enable the removal of geometric distortions induced by the GR-EPI sequence. Each run lasted 14:33 minutes, consisting of 1200 volumes with a TR of 720 ms and 2 mm isotropic voxels. Acquisitions were performed with eyes open, and relaxed fixation on a cross-hair projection over a dark background. See Ŭgurbil et al. [24] for further details on the imaging protocol.

### 2.2 Data & pre-processing

Methods pertaining to fMRI data are identical to ones the ones in Dubois et al. [1, 2] and Vieira et al. [19], and, to remove ambiguities, some steps are reprised exactly below.

Original data were provided by the HCP [25]. Preprocessed RS-fMRI data were provided by Dubois et al. [1], based on minimally preprocessed data [26]. Test scores from 1206 subjects were obtained by Dubois et al. [1]. Tests include 7 tasks from the NIH Toolbox for Assessment of Neurological and Behavioral function (NIHTB) (dimensional change card sort; flanker inhibitory control and attention; list sorting working memory; picture sequence memory; picture vocabulary; pattern comparison processing speed; oral reading recognition) and 3 from the Penn Computerized Neurocognitive Battery (Penn CNB) (Penn progressive matrices; Penn word memory test; variable short Penn line orientation). 23 subjects with missing or incomplete test scores were excluded (n = 1183). 2 subjects that scored 26 or less in the Mini Mental State Examination (MMSE) were also excluded (n = 1181). Subjects that completed four imaging sessions (n = 998) were further filtered by excluding 114 subjects with excessive in-scanner head movement (n = 884).

FMRI preprocessing included the following steps: Z-scoring normalization; removal of temporal drifts from cerebrospinal fluid (CSF) and white matter (WM); regression of mean CSF and WM signals from gray matter (GM) voxels; 12-parameter motion regression; low-pass filtering with a Gaussian kernel with a standard deviation of 720 ms; temporal drift removal from GM using a third-degree Legendre polynomial regressor; global signal regression (GSR). See Dubois et al. [1, 2] for further details. We further removed 11 subjects that had acquisitions shorter than 1200 timepoints in any RS-fMRI session, achieving a final set of 873 subjects (466 females, 407 males) with complete imaging and behavioral data. See Figure S1 for an overview of the characteristics of the final sample. Multimodal Surface Matching data were used to extract time series using the MMP-1.0 atlas [27], comprising cortical 360 ROIs, 180 in each hemisphere.

We obtained scores from 58 behavioral measures, as in Kong et al. [4] and He et al. [7]. These span domains such as alertness, cognition, emotion, motility, personality, and sensoriality (see https://wiki.humanconnectome. org/display/PublicData/HCP-YA+Data+Dictionary-+Updated+for+the+ 1200+Subject+Release). Some scores were obtained during task fMRI acquisition.

Differently from Vieira et al. [19], we did not perform band-pass filtering. There is evidence that blood-oxygen-level-dependent (BOLD) signal has higher-frequency components [28]. Besides, convolutional neural networks (CNNs) are better able to deal with the long sequences presented by high-sampled signals than RNNs.

### 2.3 Neural network architecture

We implemented an InceptionTime TCNN [23], in Flux [29], a machine learning framework in Julia [30]. InceptionTime is a 1D CNN tailored for time series classification. Based on results in Ismail Fawaz et al. [23], we opted to use 512 channels, due to the number of outputs in our networks. We also exchanged the rectified linear unit (ReLU) activation in the original for leaky ReLUs, to deal with the dead ReLU phenomenon [31]. We adapted the learning-rate decay routine in the main source repository (https://github.com/hfawaz/InceptionTime). Instead of decaying the learning-rate after only 50 epochs, we instead kept the learning-rate fixed for 800 epochs. Then, we enabled learning-rate decay, halving learning-rate after 50 epochs if training loss did not improve by a pre-fixed factor. We set the minimal reachable learning-rate to be one tenth of the original learning-rate. This learning-rate scheme should allow for an initial “exploration” of solution space. The latter phase, with smaller learning-rate, should bring the network closer to convergence. The models were trained for 1500 epochs, with a batch size of 8 subjects, which entails 32 RS-fMRI sessions per batch, using Adam [32] with an initial learning-rate of 0.001. We increased the number of networks in the ensemble to 10, from 5 in the original InceptionTime [23]. Networks are architecturally identical, but are initialized with different random seeds and trained independently. Since each participant has 4 RS-fMRI runs in the HCP, we average the prediction of each network across runs during training and testing.

### 2.4 Kernel regression

We implemented KRR as in He et al. [7]. Briefly, we trained one KRR model for each of the 58 tasks. Training includes tuning of the regularization hyperparameter, performed with an inner 3-fold cross-validation (CV). We used the same hyperparameter list of https://github.com/ ThomasYeoLab/CBIG/tree/master/stable_projects/predict_phenotypes/ He2019_KRDNN. The correlation kernel, defined as the correlation coefficient between the upper-triangular part of the average RSFC matrix between subjects, was used, as in He et al. [7].

### 2.5 Cross-validation

Outer CV was performed by splitting subjects into 20 folds, with 19 folds used for training and one fold used for validation, as in He et al. [7]. Since the HCP contains family-related subjects, we did not split families across folds, as is commonly advocated in the literature [1, 7], to avoid overestimating performance due to shared genetic and environmental factors shaping brain and behavior.

The target traits were standardized to have zero mean and unit variance according to the respective training set. This is necessary for the training of TCNNs, otherwise the loss function would be dominated by variables with higher variance. For KRR, this step is not strictly necessary, but it ensures that the hyperparameter search is consistent across traits, since the regularization depends on the scale of the target variable. Thus, instead of predicting the original target *y_i_*, we predict the standardized target 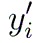, defined as:

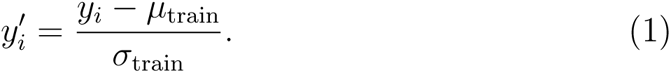

This standardization is undone to obtain predictions in the original scale to compute, *e.g.*, the mean squared error (MSE) across folds.

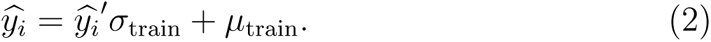

### 2.6 Performance estimates

We report Pearson’s correlation coefficient between predicted and true values, a common metric in the literature. For a given fold *f*, with *n* = *|f|* elements, predictions 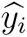 and true values *y_i_*, it is defined as

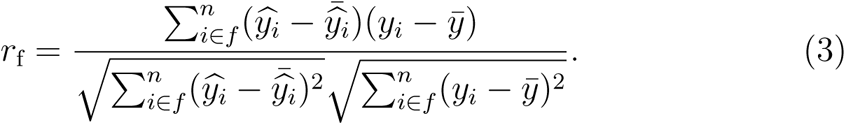

The estimate depends simultaneously on the covariance between predicted and true values, as well as the variance of each in the validation set for a given fold. The latter becomes a source of variability in the estimate, which is aggravated by small validation set sizes.

To account for this, we report the MSE across folds, which circumvents part of the excess variability. It is defined, for a given fold *f*, as

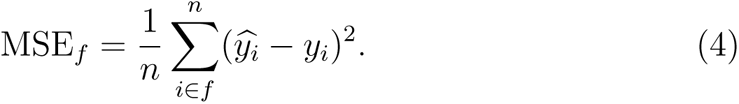

The Wilcoxon signed rank exact test was used to compare performance estimates, paired between folds. The Benjamini-Hochberg procedure was used to control the false discovery rate (FDR) among the 58 traits [33]. The false coverage-statement rate (FCR) was controlled for confidence intervals of 95% [34].

### 2.7 Post-hoc analysis: comparison with a constant model

A null level of performance cannot be directly obtained from MSE estimates, whereas for the Pearson’s correlation coefficient a value of zero indicates no correlation. For this reason, in a separate analysis, we additionally estimated a supplementary constant model, given by the mean of the true values in the training set, thus avoiding leakage, per trait. This model forfeits any information about the RS-fMRI data, and thus constitutes a ‘no-information’ model. For each significant difference in MSE between the TCNN and KRR, we performed a Wilcoxon signed rank exact test between the best performing model and the constant model.

### 2.8 Missing data

Sixteen subjects had missing data for one or more traits, and 28 traits had missing data for one or more subjects. See Figure S2 for a detailed overview. In He et al. [7], only subjects with complete behavioral assessment were used. We opted to use all available subjects, thus increasing the size of available training data. We instead used a weighted loss function to ignore missing data, in both training and assessments in validation data.

Given the individual loss, for a subject *i* and an outcome variable *j*, 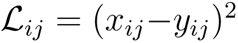, we define the weighted loss as 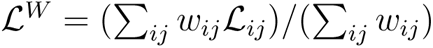. *x_ij_* is the prediction, *y_ij_* is the true value and *w_ij_ ∈* (0, 1) is the weighting, where *w_ij_* = 0 when data is missing.

### 2.9 Performance scalability

We investigated the scalability of the MSE of the TCNN and KRR models as a function of the number of subjects in the training set and the number of time points available for training.

Scalability was tested by training on subsamples, varying the number of samples available between 25%, 50% and 100% of each training fold. To ensure concordance between fractions, bigger subsamples are obtained by adding subjects to smaller subsamples, *e.g.*, the 50% subsample of a given fold is obtained by adding subjects to the 25% subsample. The number of optimization steps was kept approximately constant by increasing the number of epochs accordingly, *i.e.*, 3000 epochs for half of the subjects, 6000 epochs for a quarter of the subjects.

To test the generalizability of the models to the number of time points available during testing, we subsampled the time series to have 300, 600, 1200, 2400, or 4800 time points, respectively 3.6, 7.2, 14.4, 28.8, and 57.6 minutes. To keep this scenario realistic, accounting for effects particular to the first session [5], fractions are obtained from the earliest time points onwards. This translates the fractions into a quarter of the first session, half of the first session, the whole first session, the whole first two sessions, and all four sessions. For the TCNN, data can be used as is, since the network is fully convolutional. For the KRR, new RSFC matrices were computed from the available time points, with averaging of the RSFC matrices across sessions when necessary, as performed in the training procedure.

### 2.10 Confounder variables

In He et al. [7], deconfounding was performed in training data with a linear model and applied to validation data. This choice, albeit justified, negatively biases results and does not guarantee prediction and confounder to be uncorrelated across test folds [35]. An alternative based on the removal of linear effects of confounders from features can lead to leakage [36]. The approach presented in Chyzhyk et al. [35], anti mutual-information subsampling, is a possible solution to these issues but cannot be easily adapted to our problem. Since we have 58 output variables at once, forming test samples that optimize anti mutual-information becomes a harder problem.

In Dubois et al. [2] authors predicted personality factors and fluid intelligence from RSFC, and controlled for several confounders, *i.e.* sex, age, fluid intelligence, brain size, in-scanner movement and software reconstruction version. However, they argue that some of these variables that could be seen as confounders can also be causally linked to the variables of interest, possibly leading to “pessimistic” performance estimates. In our case, given that the variables of interest include domains beyond personality, to include even motility and sensoriality, even “obvious” confounders such as in-scanner movement can display causal relationships with some of the traits.

We instead opted to not perform covariate regression during training. We report results without covariate regression and with a post-hoc confounder regression. This procedure consists of estimating a linear model that predicts the outputs from confounders variables in training data, and using this regression to subtract the effect of confounders from validation data. We considered gender, age, brain volume, movement from each resting-state session, and reconstruction algorithm version as confounders, as in Dubois et al. [1] and Vieira et al. [19].

### 2.11 Regional importance

We investigated the importance of each region to the predictions of the TCNN models by computing saliencies in the validation data, i.e. the gradient of the predicted value with respect to the input time series, as in Vieira et al. [19]. This is a measure of how much the prediction would change if the time series of a given region was changed. Since we account for the fact that the timeseries are standardized, resulting saliencies have mean zero. We extracted the mean squared saliency for each region, and averaged across subjects in the same fold. We then computed the rank of the mean squared saliency across regions, and averaged across folds, resulting in a mean rank for each region. Due to the large rank correlation between regions, we report the mean rank across all regions.

## 3 Results

Several traits were predicted with Pearson’s correlation coefficient between estimated and true values significantly different from zero. Both the TCNN and KRR could predict 41 and 42 traits, respectively, with a significant correlation coefficient, without covariate regression, with 40 traits in common. Both models could predict the five NEO-Five Factor Inventory (NEOFFI) personality traits, with performance approaching *r ≈* 0.2 for NEOFAC C and *r ≈* 0.3 for NEOFAC O. Performance is also significantly different from zero for eleven cognitive scores, including *r ≈* 0.4 for ReadEng Unadj and PicVocab Unadj, and *r ≈* 0.3 for PMAT24 A CR and VSPLOT TC. However, TCNN performance for Emotion Task Face Acc and ER40FEAR, and the KRR performance for PercReject Unadj were not significantly different from zero, after FCR correction.

Figures 1a and 1b illustrate the CV Pearson’s correlation coefficients obtained, without and with deconfounding. Deconfounding regression was always estimated on training data and applied to validation data. Shown are the expected values, with ellipses highlighting the 0.995 confidence interval of the mean, based on a bivariate t-student distribution. In opposition to He et al. [7], we did not find significant differences in the Pearson’s correlation coefficients between predicted and true target variables after correcting for multiple comparisons. The mean Pearson’s correlation coefficients and confidence intervals for all traits are shown in Figures S3 and S4, including FCR correction [34]. For all results, we removed performance estimates from four folds in the Mars Final Contrast Sensitivity

**Figure 1:**
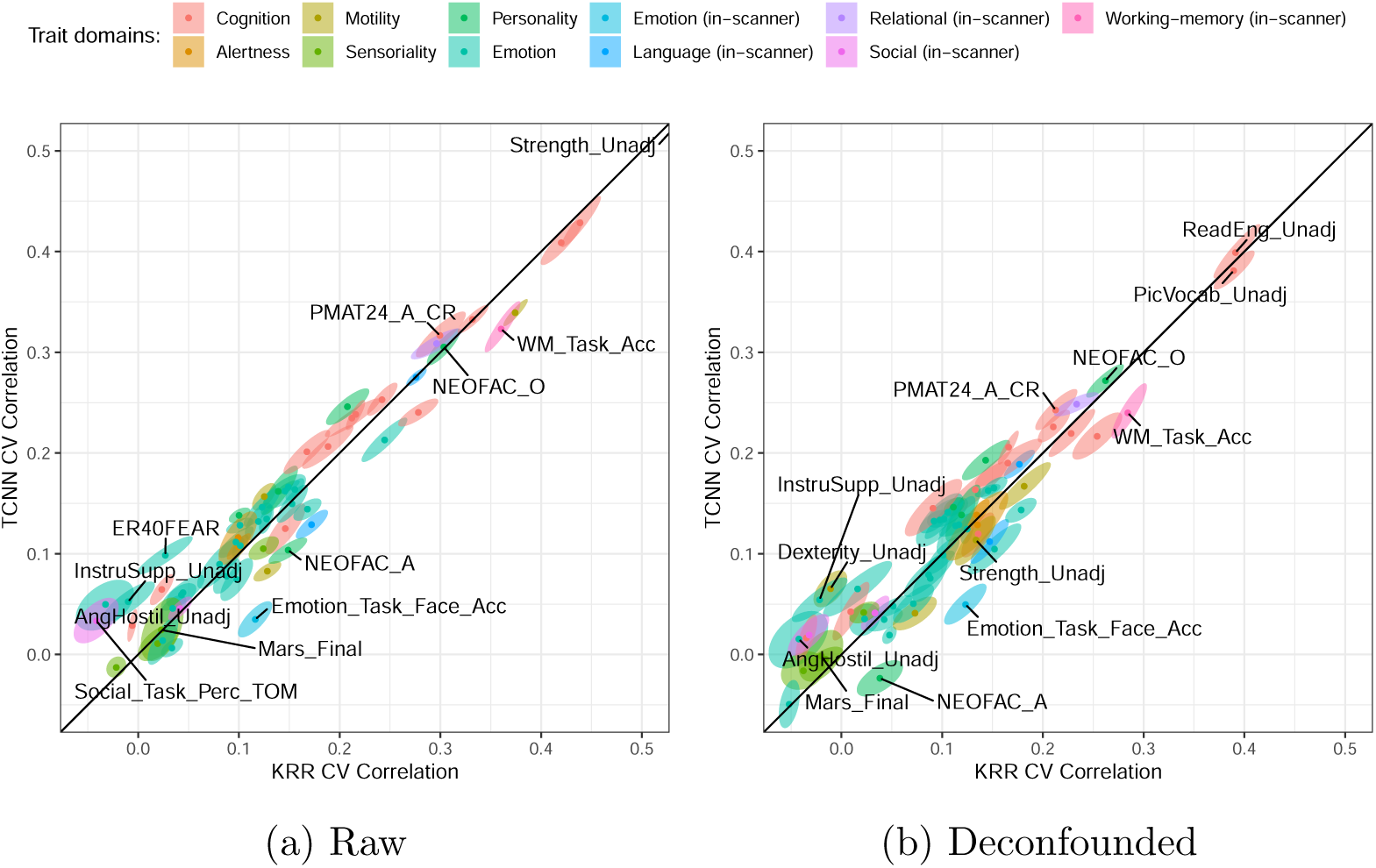
CV results in terms of Pearson’s correlation coefficient. (a) shows results without deconfounding while (b) includes deconfounding of predicted values and true labels. Ellipses delineate the 0.995 confidence region for the average based on bivariate t-statistics, not corrected for FCR. Domains are given by the HCP data dictionary.

Score (“Mars Final”) where MSEs clearly displayed outlier behavior for both the TCNN and KRR, being orders of magnitude greater than other folds. This happened due to chance in how subjects were allocated to folds, and the fact that both the evaluation and training objectives are highly sensitive to outliers. The removal did not change significance, but altered the ranking of this trait in different analyses.

Figure 2 shows the difference in MSE between the TCNN and KRR models, taking into account covariate regression. Only traits with a significant difference between models are shown, including FDR-correction for multiple comparisons. Eleven significant differences between the models were found in terms of raw MSE. In seven, the TCNN achieved lower MSE than KRR. These include four Penn Emotion Recognition Test (ER-40) scores: Number of Correct Sad Identifications (“ER40SAD”), Fear Identifications (“ER40FEAR”), Anger Identifications (“ER40ANG”), and Happy Identifications (“ER40HAP”). Additionally, the “Overall Percentage of stimuli that the subject rated as ‘random’ in SOCIAL task” (“Social Task Perc Random”), which is a score from a task performed during a separate fMRI acquisition, the Mars Final Contrast Sensitivity Score (“Mars Final”), and the NEOFFI Conscientiousness Scale Score (“NEOFAC C”), were also better predicted by the TCNN. On the other hand, the KRR better predicted the “Accuracy Percentage during FACE blocks in EMOTION task” (“Emotion Task Face Acc”), also an in-scanner task from a separate fMRI acquisition, the NEOFFI Agreeableness Scale Score (“NEOFAC A”), NIHTB Oral Reading Recognition Test: Unadjusted Scale Score (“ReadEng Unadj”), and the NIHTB Grip Strength Test: Unadjusted Scale Score (“Strength Unadj”). These are shown in Figure 2a. When performing post-hoc covariate regression, the number of significant differences increased to 13. In this regime, the TCNN was able to predict 11 traits with lower MSE than KRR, still including several ER-40 scores, while the KRR better predicted two traits. For the full list of traits, see Tables S2 and S3.

**Figure 2:**
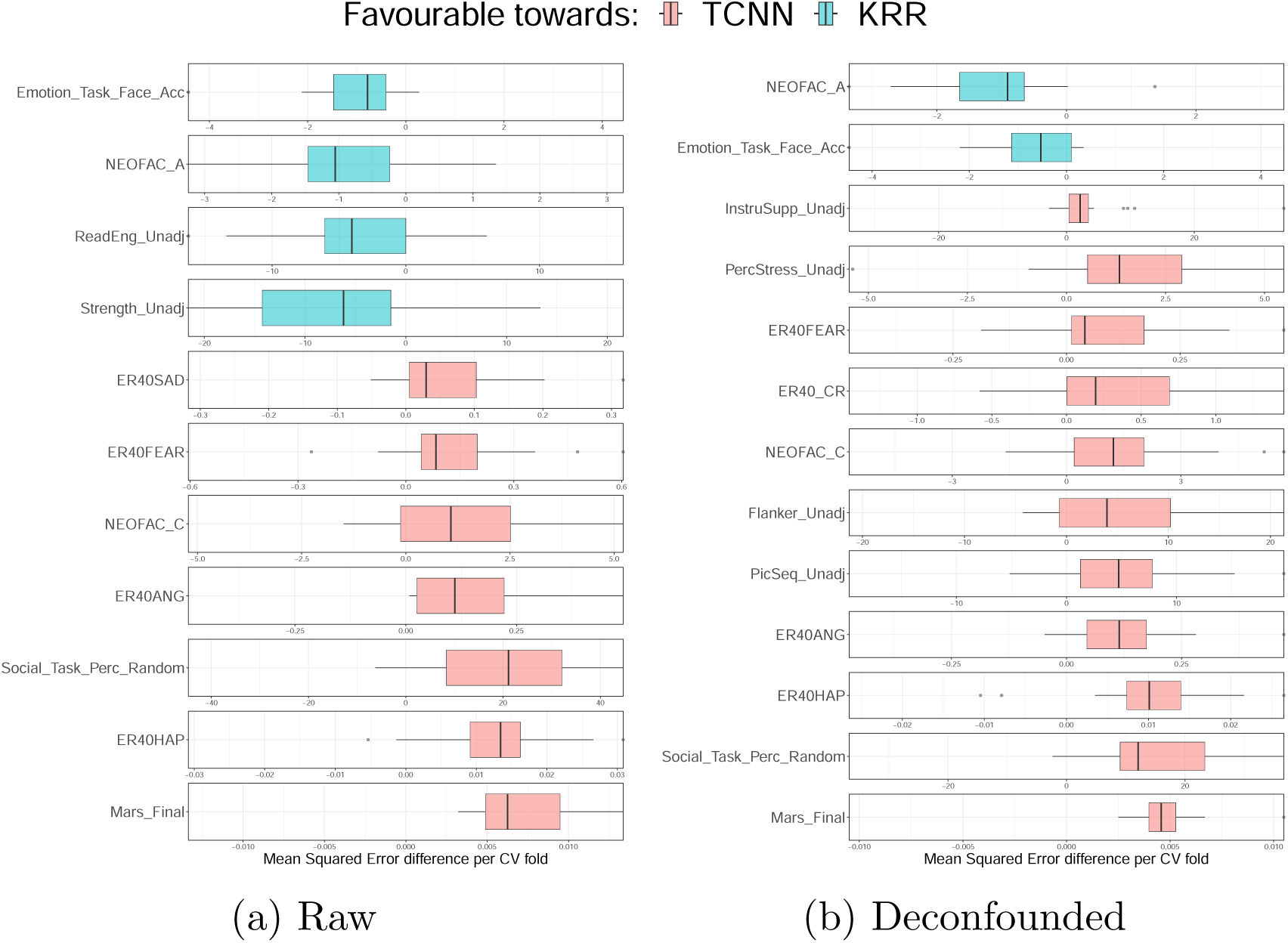
CV results in terms of paired MSE differences without covariate regression. (a) and after post-hoc covariate regression (b). Performance differences are shown in the original scale. Due to the different scales between traits, and to make visualization easier, we center the plots around zero, which highlights the direction, magnitude and variability of the differences. Only traits with significant differences (*p <* 0.05, corrected for multiple comparisons) are shown. Significance was assessed by a Wilcoxon signed rank exact test.

Post-hoc testing with a constant model showed that, when not accounting for covariates (Table S4), performance of the TCNN were significantly better than the constant model for “NEOFAC C”, while the KRR was significantly better for “Emotion Task Face Acc”, “ReadEng Unadj” and “Strength Unadj”. If accounting for covariates (Table S5), the TCNN was significantly better than the constant model in all the comparisons shown in Figure 2b, whereas the KRR was significantly better for “NEOFAC A”.

To provide an illustrative comparison between models’ performance for future studies, Figure 3 shows the actual MSE distributions for the same traits in Figure 2. Figure 3a shows the raw MSE distributions, while Figure 3b shows the distributions after post-hoc covariate regression. Outliers are clearly visible in the results, coming from folds having much larger MSE values. The four extreme outliers from the Mars Final Contrast Sensitivity Score are again not shown.

**Figure 3:**
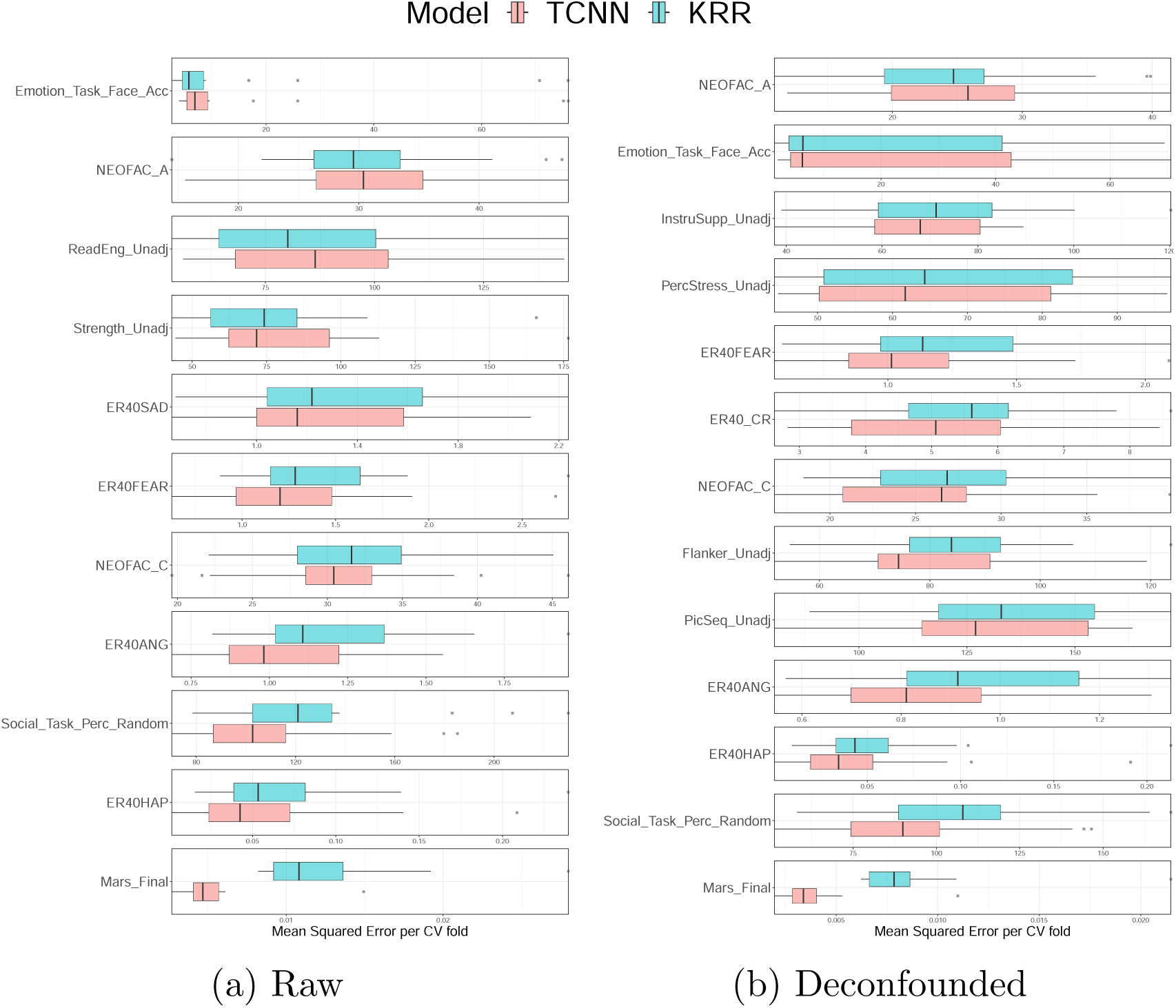
CV results in terms of MSE, without covariate regression. (a) and after post-hoc covariate regression (b). Only traits with significant differences (*p <* 0.05, corrected for multiple comparisons) are shown. Significance was assessed by a Wilcoxon signed rank exact test.

When using the Pearson’s correlation coefficient as the performance metric, no difference in performance scaling was detected for any trait with the number of subjects in the training set or the number of time points available during testing. On the other hand, with MSE as the performance metric, significant differences were found in the scaling with the number of subjects in the training set and the number of time points available during testing. Out of 18 significant differences, in 17 traits increasing the number of subjects resulted in the MSE of KRR progressively approaching that of the TCNN (Figure S5). From the 36 identified differences, in 32 traits increasing the number of time points resulted in the MSE of TCNN progressively approaching that of the KRR (Figure S6). Or, in other words, the KRR was more sensitive to the variation in the number of subjects in the training set, while the TCNN performance was more affected by the number of time points available during testing. See Figures S7 to S10 for the respective scaling curves.

The average ranks of the mean squared saliency of the TCNNs, averaged across folds and traits, are shown in Figure 4, generated with ciftiTools [37]. The correlation of the mean squared saliency between traits is very high, exceeding *r* = 0.9 for any pair of traits.

**Figure 4:**
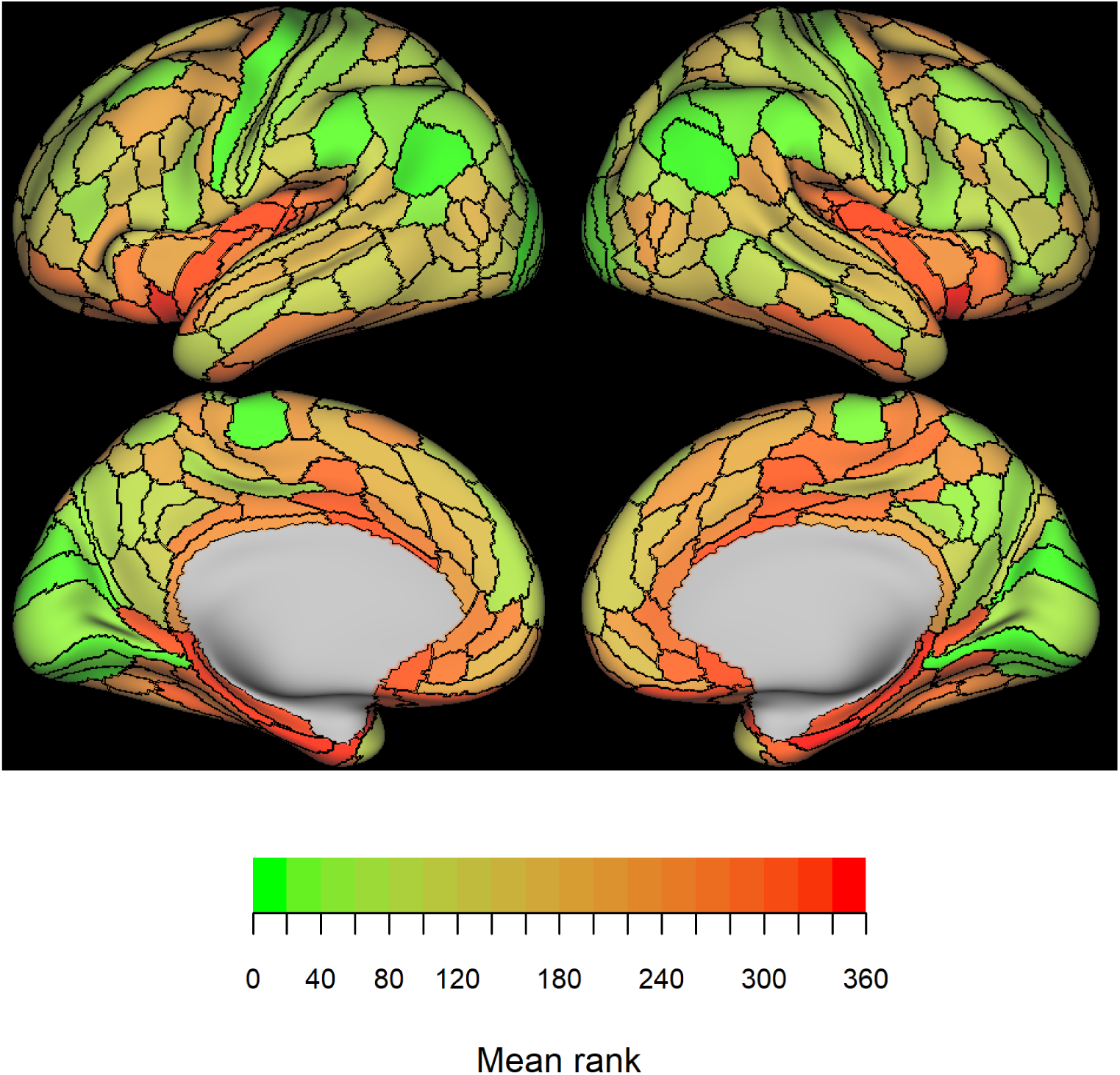
Average rank of the squared sum of saliencies of the TCNNs. For each trait, the squared saliency was averaged across subjects per fold, and the rank of the mean squared saliency was computed across regions. Ranks were then averaged across folds and traits.

## 4 Discussion & Conclusion

We demonstrated that the majority of 58 traits in the HCP can be predicted from RS-fMRI data with timeseries-based TCNN models with performance significantly different from zero. These span domains such as cognition, motility, personality, emotion and even performance in tests performed during separate task-fMRI sessions. Compared to a strong RSFC-based baseline, KRR, we found that the TCNN performs significantly better in a number of traits.

Well performing traits include nine out of ten test scores in the general intelligence factor formulation in Dubois et al. [1]. The exception is the Penn CNB Word Memory Test: Total Number of Correct Responses (“IWRD TOT”), both when regression of covariates is performed and when it is not (see Figures S3 and S4). This test score achieved non-significant performance in previous studies [6]. Picture Vocabulary (“PicVocab Unadj”) and Oral Reading Recognition (“ReadEng Unadj”) are highlighted in Kashyap et al. [5], Li et al. [6], He et al. [7], and Kong et al. [11] as being the best predicted items. Their results included covariate regression, and likewise we find that these two items are the best predicted traits in our analogous analysis (Figure 1b). These two items comprise the crystallized ability factor in Dubois et al. [1], and heavily load on the general factor^1^. Likewise, the Penn Progressive Matrices: Number of Correct Responses (“PMAT24 A CR”) is predicted with a smaller, but still substantial correlation, at a level comparable to the literature [2, 3, 7, 17, 38], and also highly loads on the general factor [1]. This provides evidence that the prediction of specific traits aligns with the prediction of the general factor, with implications for the study of the neural correlates of cognition.

Personality traits based on the NEOFFI scores are also predicted with a significant correlation coefficient by both models. Our results agree with Dubois et al. [2], including the lack of significant correlation for Agreeableness (“NEOFAC A”). Both models were able to predict Openness to experience (“NEOFAC O”), with performance similar to *r* = 0.24 reported in Dubois et al. [2]. The TCNN achieved significantly lower MSE than KRR in the prediction of Conscientiousness “NEOFAC C”, the second best predicted NEOFFI trait in Dubois et al. [2], as well as in our results.

Several ER-40 scores were also better predicted by the TCNN in Figures 2 and 3. These include the Number of Correct Happy Identifications (“ER40HAP”), Anger Identifications (“ER40ANG”), Fear Identifications (“ER40FEAR”) and the Number of Correct Responses (“ER40 CR”). The latter has been shown to be significantly, albeit weakly, associated with schizophrenic or schizoaffective diagnoses [39, 40]. One possible explanation for better performance of the TCNN in Emotion Recognition is that neuronal correlates of ER-40 scores are manifest in the RS-fMRI data as temporal patterns, such as phase differences or waveforms, which are better captured by the TCNN than by KRR. However, the TCNN is still not able to predict the ER-40 scores with high accuracy, with *r ≈* 0.1 for the best predicted trait, “ER40FEAR”. In a post-hoc analysis with an estimated constant model, we found that when accounting for covariates the TCNN was significantly better than the constant model for all aforementioned ER-40 scores (Table S5), although it does not differ significantly from the constant model if covariates are not accounted for (Table S4). This reiterates the very low effect size of machine learning model performance in the prediction of ER-40 scores, which can also be attested in the literature [6].

We show that, as expected, covariate regression directly affects performance estimates. While within individual folds performance might change positively or negatively, the overall effect is a decrease in Pearson’s correlation coefficients following removal of covariate effects (Figure S11). This phenomenon occurs across both models. Grip strength (“Strength – Unadj”) exhibits an interesting pattern, also visible in Figure 1. It demonstrates the highest correlation among traits for both models, in excess of *r >* 0.7 when not performing covariate regression. However, when removing possible confounding effects, the correlation, although still significant, is greatly diminished to *r ≈* 0.1. The level of performance obtained after deconfounding is compatible with Kashyap et al. [5] (*r ≈* (0.1, 0.2)), Kong et al. [4] (*r ≈* (0.06, 0.14)), Li et al. [6] (*r ≈* (0.15, 0.23) with and without GSR), He et al. [7] (*r ≈* (0.13, 0.2) with KRR only, while fully-connected neural network (FNN), BrainNetCNN and Graph CNN (GCNN) heavily underperform), Kong et al. [11] (*r ≈* (0.1, 0.16)). Grip strength is strongly correlated (*r* = 0.3659) [6] with in-scanner head motion in the HCP, measured by DVARS, with known systematic effects on fMRI data [41]. Additionally, grip strength is strongly correlated with sex [42], which can be predicted from RS-fMRI data with accuracy surpassing 90% [20]. We can then conclude that high performance on the prediction of grip strength is related to that of covariates, sharing information in the temporal dynamics of RS-fMRI data, as well as in RSFC.

Performance is highly correlated between models, within and between traits, as can be visually assessed in Figure 1. This hints towards a common source of information, *i.e.* features extracted by the TCNN span approximately the same manifold as those extracted by KRR. See Figures S3 and S4 for the full list of performance estimates. Using the same categorization from Li et al. [6] and Líegeois et al. [43], Figure S12 demonstrates that the highest performance for the timeseries-based TCNN model is also achieved in traits that are derived from tasks, instead of self-report questionnaires, further corroborating the similarity between models’ performances.

We did not find any significant differences in Pearson’s correlation coefficients between the two models after correcting for multiple comparisons. This is in contrast to the findings of He et al. [7], who found two significant differences in Pearson’s correlation coefficients between models. In their results, KRR predicts grip strength (“Strength Unadj”) better than the GCNN. Also, KRR predicts picture matching vocabulary (“PicVocab – Unadj”) better than the FNN. We believe that this difference can be attributed to the fact that the Pearson’s correlation coefficient displays higher variance than the MSE. This is the case because the Pearson’s correlation coefficient is the ratio of a covariance and two standard deviations. Ratios can be thus less precisely estimated than the MSE for small samples, since they are subject to variance in the denominator. This same argument can be applied to the coefficient of determination, or R-squared, since it involves the variance of the response variable in the denominator.

One of the limiting factors to train machine learning models for the study of brain-behavior relationships is the availability of data. This is particularly true for fMRI data, which is expensive to acquire and process, and deep neural networks, which require large amounts of data to train due to their high flexibility. We were able to demonstrate the dependence of MSE on the number of subjects in the training set (see Figures S5 and S9). Overall, doubling the number of subjects in the training set results in lower MSE for both models in most traits. Against the usual expectation, however, for many traits the TCNN is able to achieve lower MSE than KRR when using only half or a quarter of the subjects in the training set (Figure S9). The only trait that does not display this behavior is the NIHTB Grip Strength Test: Unadjusted Scale Score (“Strength Unadj”). The fact that the TCNN predicts the 58 traits at once can be a possible explanation for this observation, since the joint prediction has been shown to be beneficial [44].

We also detected associations between MSE and RS-fMRI time series length for either model in a number of traits (Figures S6 and S10), particularly evident in cognitive test scores. While this is not surprising, it reaffirms that slow temporal dynamics are important for prediction of these traits. Session length dependence was also found in Jiang et al. [45] for functional connectivity-based prediction of behavior from eight fMRI tasks in the HCP, including RS-fMRI.

The performance estimates based on the Pearson’s correlation coefficient, however, do not display either phenomena (see Figure S7 for scaling on the number of subjects and Figure S8 for scaling on time series length). This underscores the importance of the choice of performance metric. We cannot exclude the possibility that previous studies that used the Pearson’s correlation coefficient as the performance metric may have missed significant differences in performance between models. While the Pearson’s correlation coefficient is a common metric in the literature, it is not without its limitations. It is location and scale invariant, as long as the rescaling is positive. While this is a desirable property when comparing across different target variables, it also means that arbitrary linear transformations of the predictions can achieve the same correlation coefficient. In other words, we can construct arbitrarily bad predictions that still achieve a high correlation coefficient.

Due to the shared information within model predictions, the saliency of the TCNN is highly correlated between traits. This is expected, since the TCNN is trained to predict all traits at once and, due to regularization, traits that are easier to predict will steer the model parameters towards their own prediction. In our case, cognitive test scores are the best predicted traits. Unsurprisingly, the saliency seen in Figure 4 shares many similarities with the one observed in Vieira et al. [19], where a RNN was used to predict general intelligence in the HCP. Highly salient regions are distributed across the brain, including multiple occipito-parietal regions, visual cortices and the primary motor cortex. On the other hand, regions with low saliency are concentrated in the insula and medial temporal cortices, including the piriform, entorhinal, perirhinal ectorhinal, anterior agranular insula, para-insular and pre-subiculum cortices, as defined in Glasser et al. [27]. These regions displayed low contrast between task fMRI conditions and, although neurobiological interpretation cannot be discarded, medial regions are known to be more susceptible to signal to noise ratio issues [27], which could attenuate the reliance of the TCNN on these regions.

Since we excluded subjects with incomplete RS-fMRI runs, and also subjects with excessive in-scanner head motion, we retrieved data from fewer subjects than He et al. [7], which excluded subjects with missing behavioral data. This poses a risk of selection bias in results, as subjects with missing behavioral data or censored subjects may present systematic differences from those with complete data. These choices were made to ensure that the RS-fMRI data is of sufficient quality.

There is a risk of overfitting the HCP data, in that newer state-of-the-art accuracies can be due to chance and not actual improvement. Prediction of behavior traits from HCP subjects has been performed in numerous studies [4–11]. Often, studies use these tasks to benchmark machine learning models. This form of validation set overfitting is notorious in computer vision benchmarks.

There is a risk that the results for KRR are optimistic in our study. KRR was shown to be a strong predictor in HCP data in multiple prior studies [4, 6, 7, 11]. This implies a level of leakage, since we are using information from the whole data to base our comparisons. This risk, however, is minimized, since we followed the exact same modelling procedure described in He et al. [7]. Additionally, the RS-fMRI pre-processing was optimized in Dubois et al. [2] for RSFC based predictive modeling. This means that the features used for the KRR were optimized based off this very same dataset, which can make a difference in comparisons. Dubois et al. [2] performed prediction of personality factors and fluid intelligence. NEOFFI scores for Conscientiousness (“NEOFAC C”) and Agreeableness (“NEOFAC A”) displayed significant differences between methods in Figures 2 and 3. Conscientiousness was better predicted by the TCNN, while Agreeableness was better predicted by KRR. We did not observe differences in fluid intelligence (“PMAT24 A CR”).

While robust strategies exist to diminish the effect of confounders, in the absence of a scalable method that could deal with the number of traits we studied in this work we used a post-hoc strategy. While typically confounders should be controlled, typical “confounder variables” can actually be part of complex causal chains in brain-behavior relationships. Since this causality is not always obvious, removing the effect of a “confounder” that mediates or moderates the brain phenotype under study would lead to spurious results. For example, age is associated with differences in several behavioral traits, and is also associated with brain phenotypes, such as cortical thinning [46] or changes in RSFC [47]. In a causal chain, brain phenotypes, of which some are unobserved or unobservable, mediate the effect of aging on behavior, since behavior is caused by the brain. Removing the effect of age would lead to spurious results, since the effect of brain phenotypes on behavior would be partially removed as well. See Smith et al. [48] for a discussion on this topic.

Future work could demonstrate if with increased sample sizes and datasets enriched with other fMRI tasks CNNs and other neural networks can surpass traditional machine learning methods. The specificity and reliability of traits and imaging measurements can bias performance estimates, *e.g.*, artefactual relationships, noisy measurements. Enhanced imaging and behavioral methods can be used, leading to improvements in prediction. Future work can also investigate the definition and role of confounder variables and how to best deal with them in prediction of behavioral traits.

Generalizable prediction of behavior from brain imaging is a challenging task. Li et al. [12] shows that for several traits included in our work, models trained on white Americans do not generalize to African Americans, in both the HCP and Adolescent Brain Cognitive Development study. Systematic biases in the data can lead to spurious results. For example, in our sample, males are substantially younger (see Figure S1), which can bias results due to an interaction between sex and age. Further research is needed to understand how to best deal with these issues, and to ensure that models generalize to diverse populations.

In conclusion, prediction of behavior based on kernel learning with RSFC or TCNN with RS-fMRI time series are comparable in terms of performance. In most traits, effect sizes are low for either model, except for cognitive test scores. This has important implications for the neurobiological basis of different dimensions of human behavior, in terms of temporal dynamics of BOLD fluctuations. However, our systematic investigation showed that differences in performance between models do exist. For certain traits and under certain conditions, a CNN-based modelling can learn more discriminative features than traditional kernel learning based on RSFC, even though TCNNs are notoriously more complex to train.

## Acknowledgements

Authors acknowledge Julien Dubois et al. for generous sharing of code and pre-processed data.

Data were provided [in part] by the Human Connectome Project, WUMinn Consortium (Principal Investigators: David Van Essen and Kamil Ugurbil; 1U54MH091657) funded by the 16 NIH Institutes and Centers that support the NIH Blueprint for Neuroscience Research; and by the McDonnell Center for Systems Neuroscience at Washington University.

Authors acknowledge funding received through grant #2018/11881-1, São Paulo Research Foundation (FAPESP) (B.H.V – PhD scholarship) and [10001C 197480], Swiss National Science Foundation at the University of Zurich (B.H.V. – postdoctoral position).

Authors acknowledge the Julia machine learning community for fruitful discussions.

## A Sample characteristics

**Figure S1:**
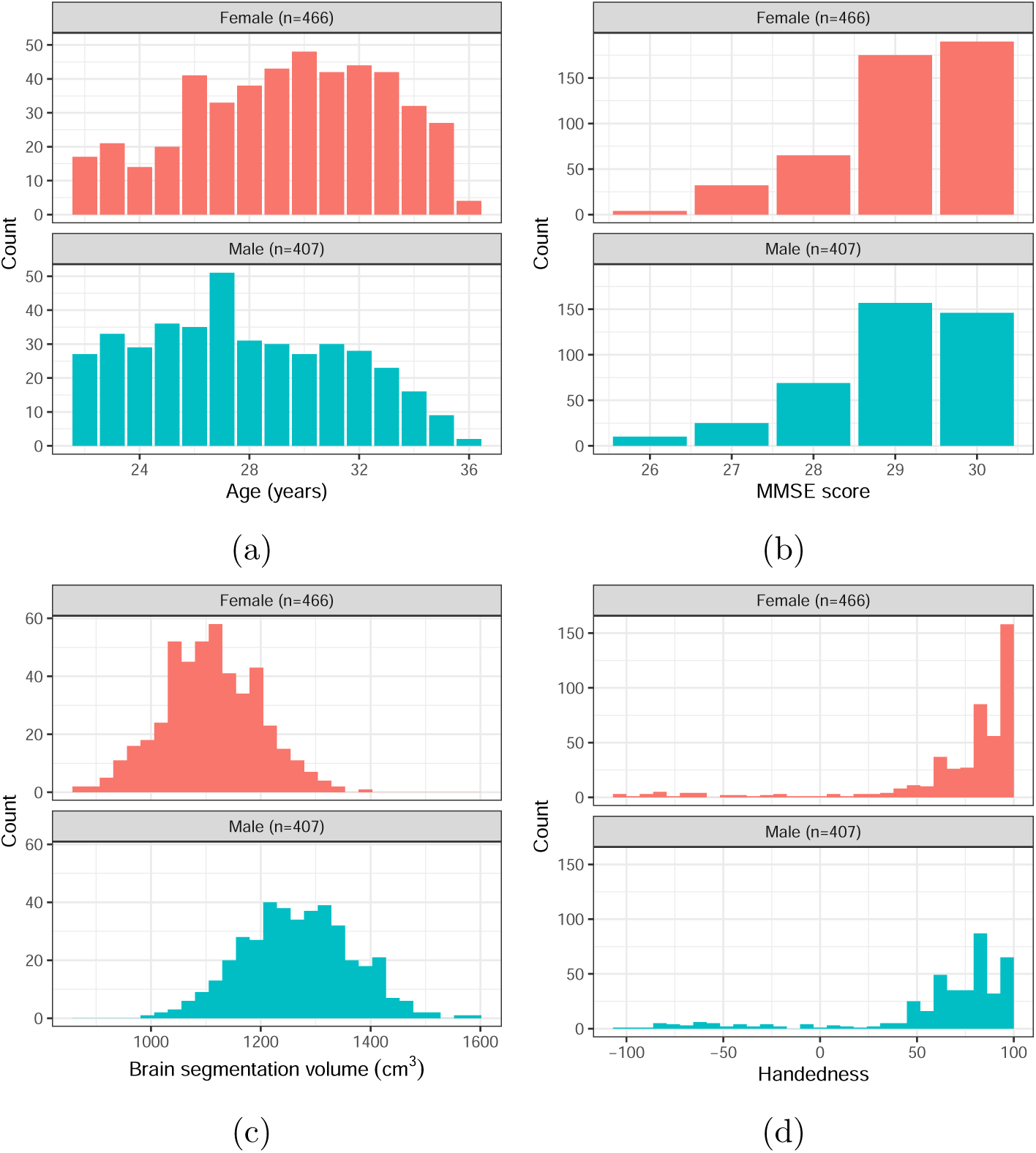
Sample characteristics of the HCP subjects included in this study, per sex. (a) Age distribution. (b) MMSE distribution. (c) Brain volume distribution. (d) Handedness distribution.

## B 58 behavior variables defined in the Human Connectome Project

**Table S1:**
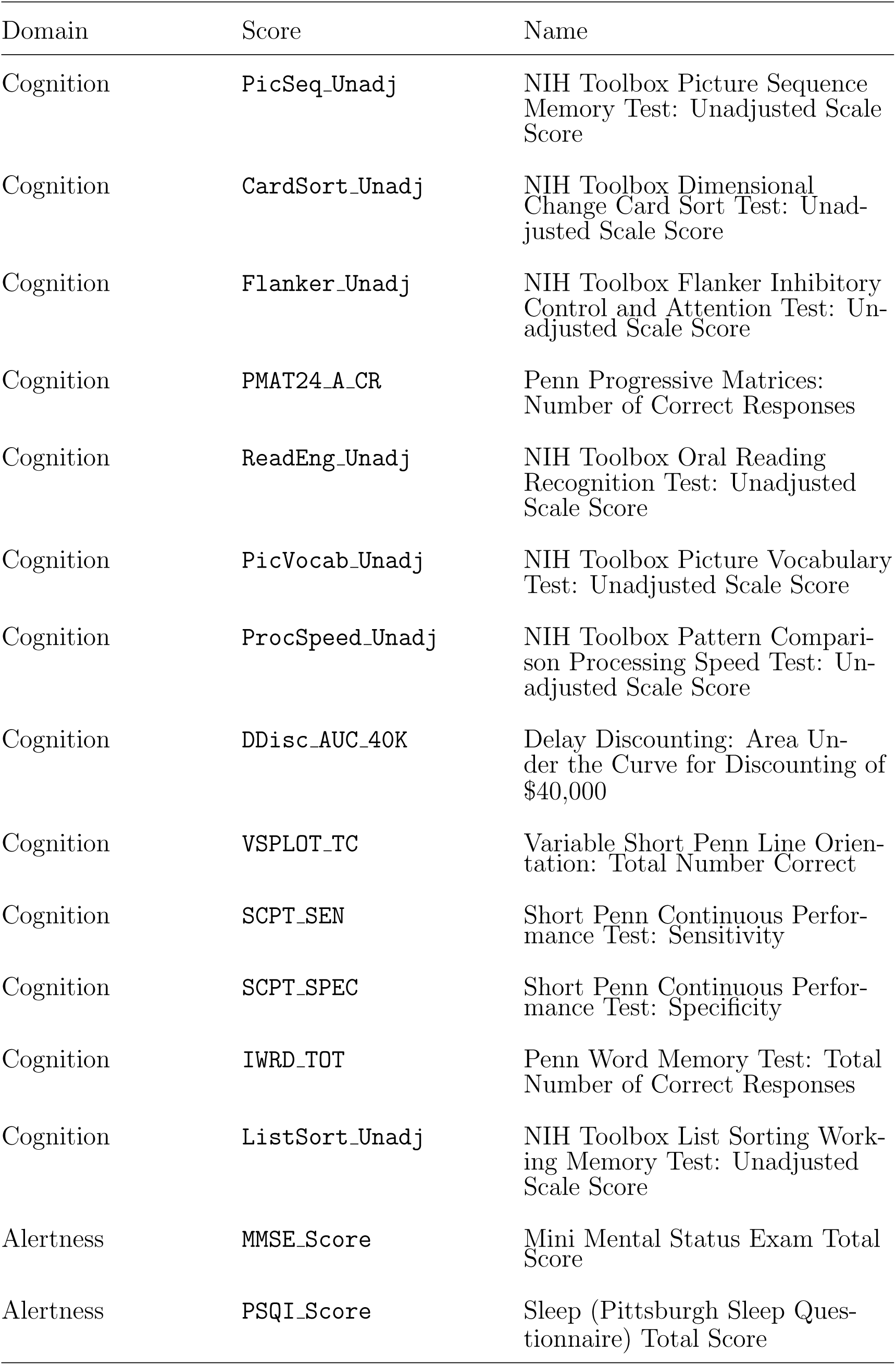
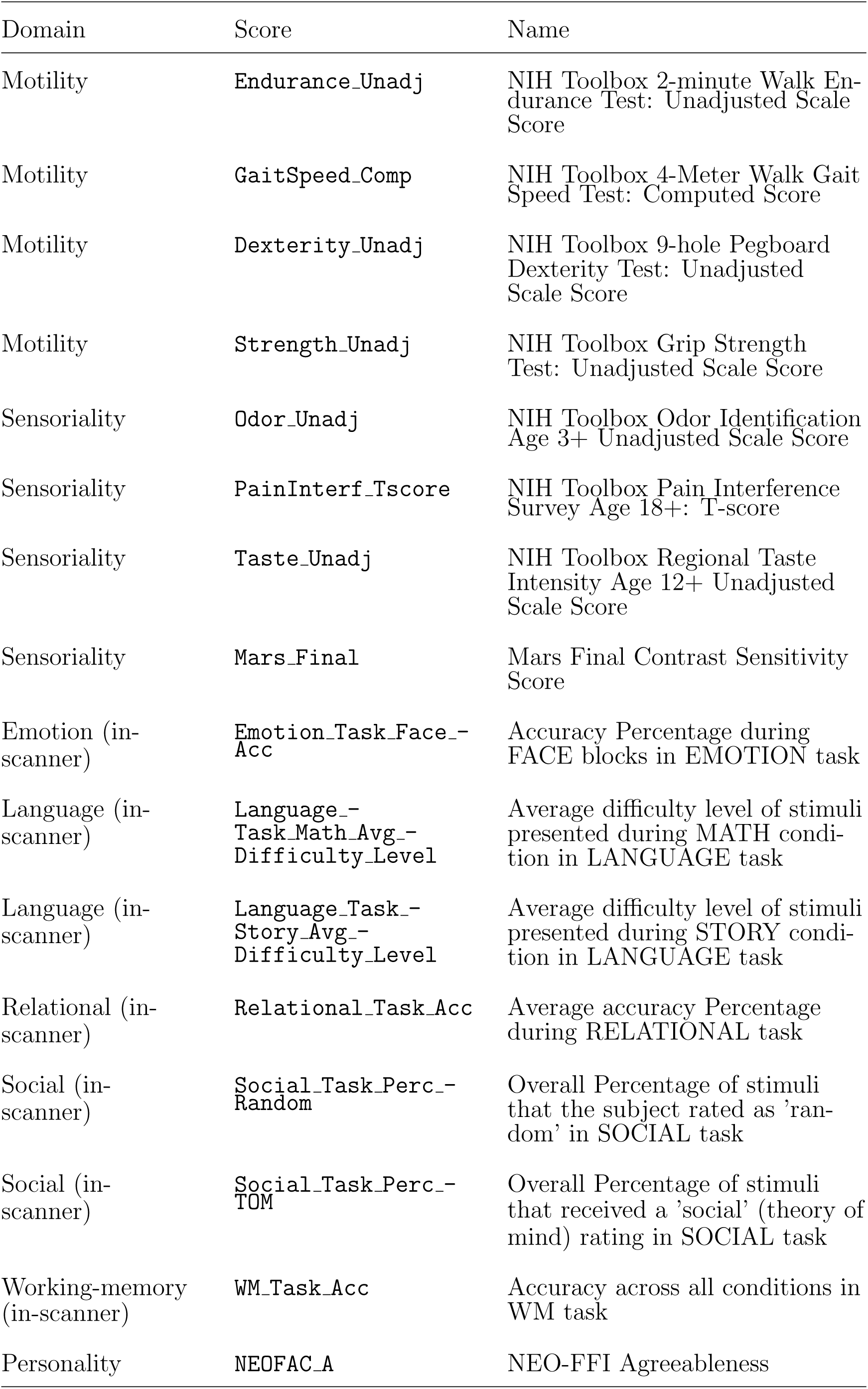
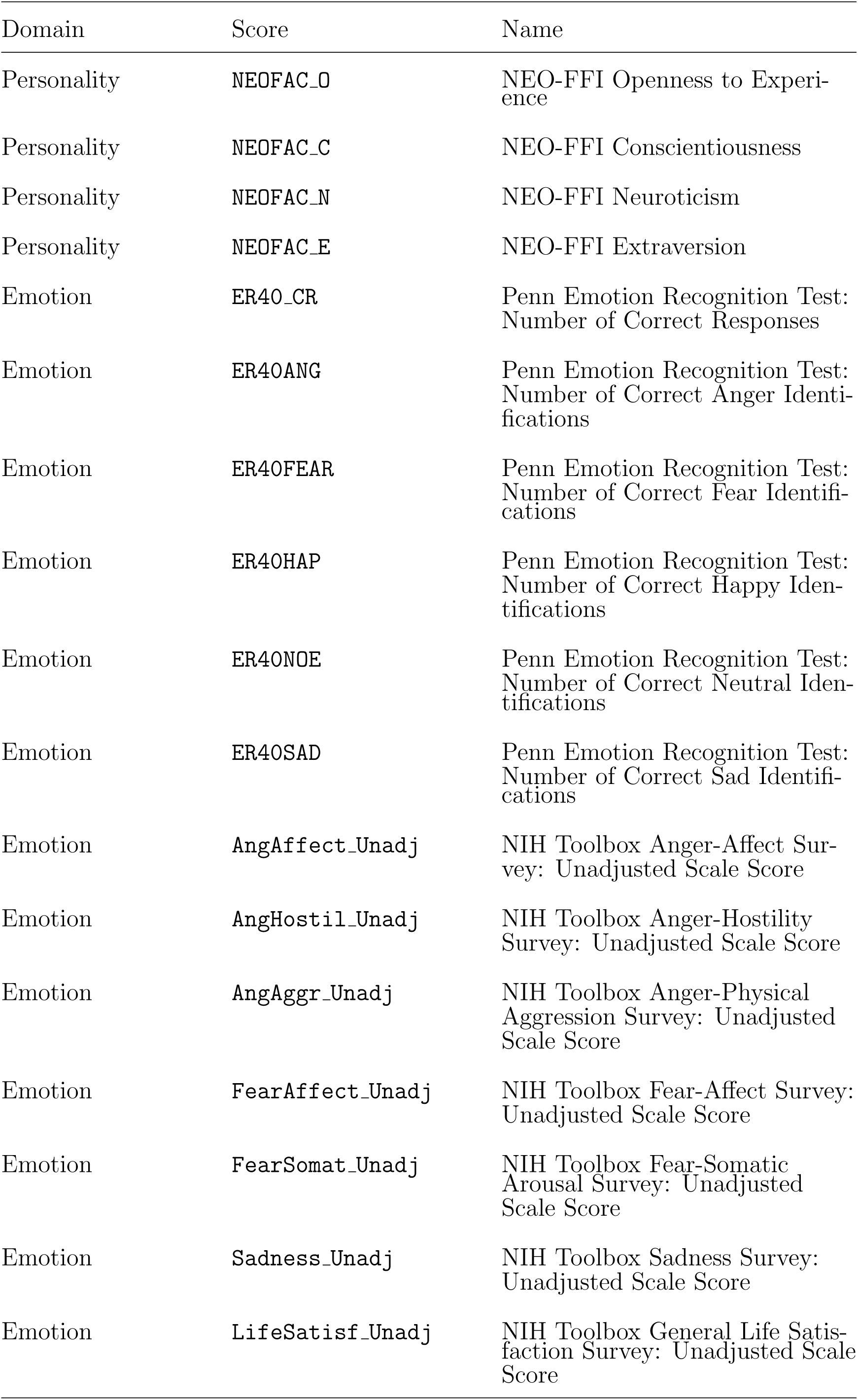
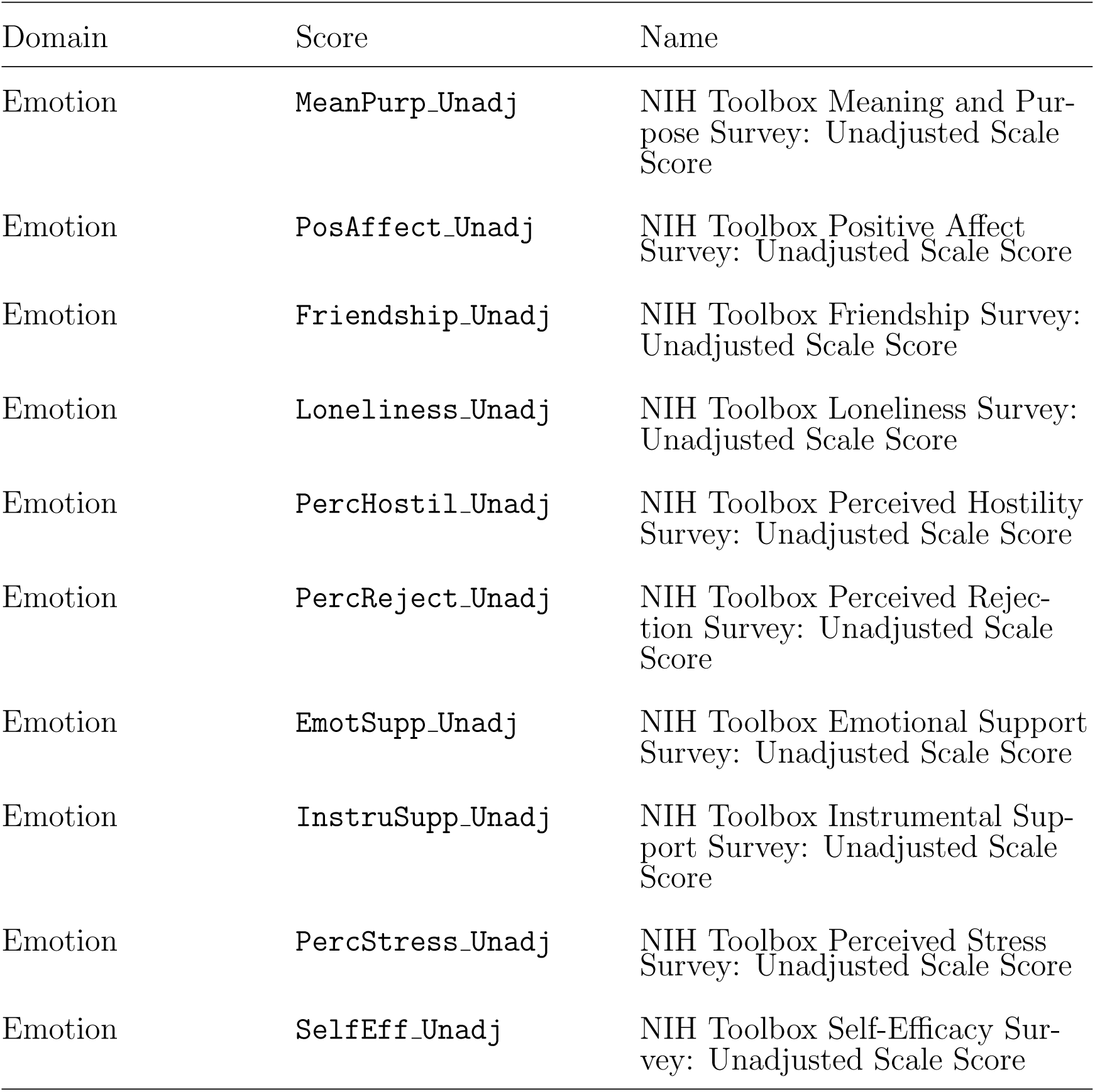
58 behavioral traits in the HCP dataset.

## C Missing data statistics

**Figure S2:**
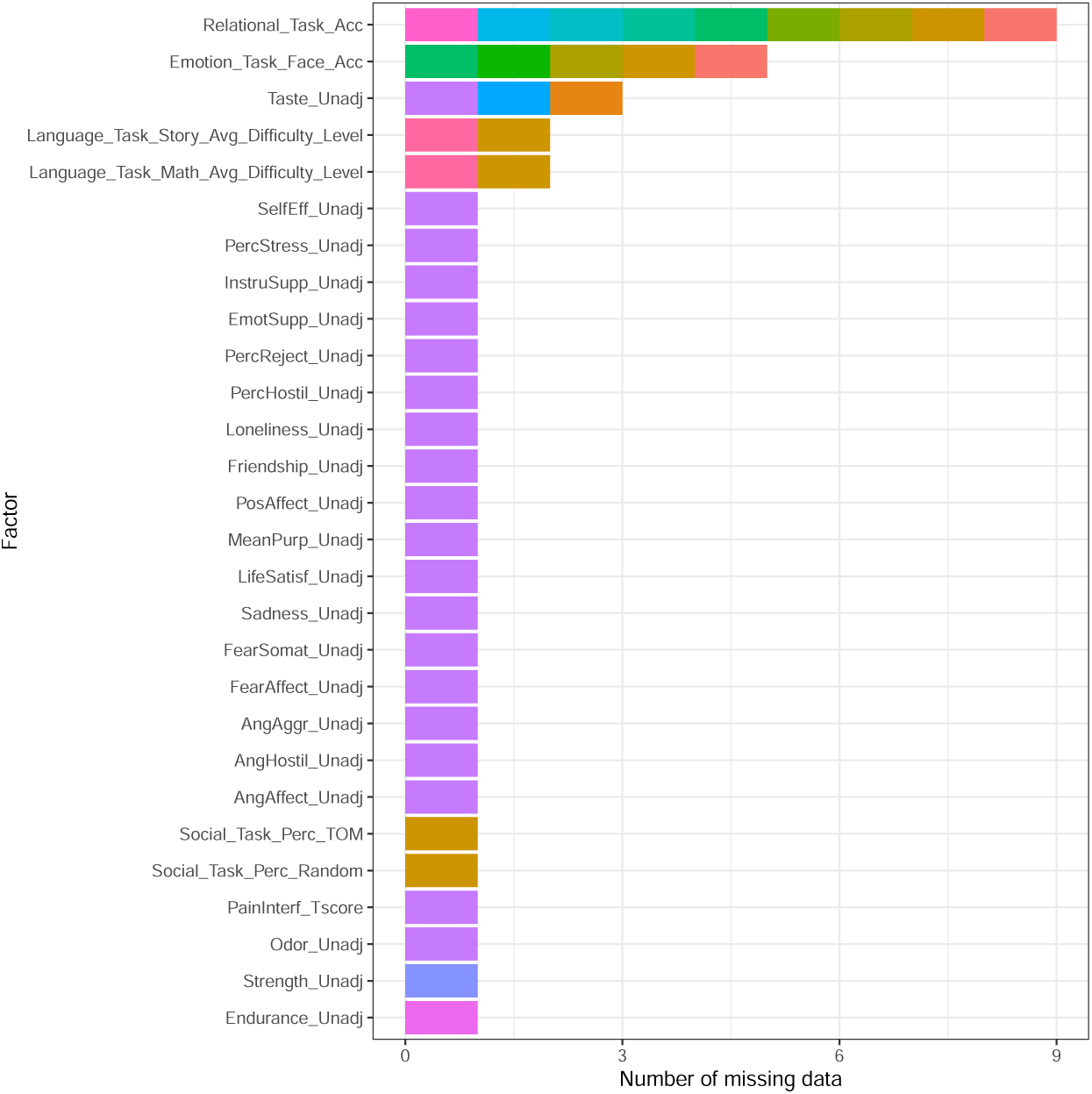
Number of subjects with missing data for each trait, for the whole sample (n=873). Each subject is represented individually by a color. Sixteen participants have missing data, for at least one trait: 12 for 1 trait, 4 for 2 traits, 1 for 6 traits, and 1 for 20 traits.

## D Prediction accuracy across 58 traits from the Human Connectome Project

**Figure S3:**
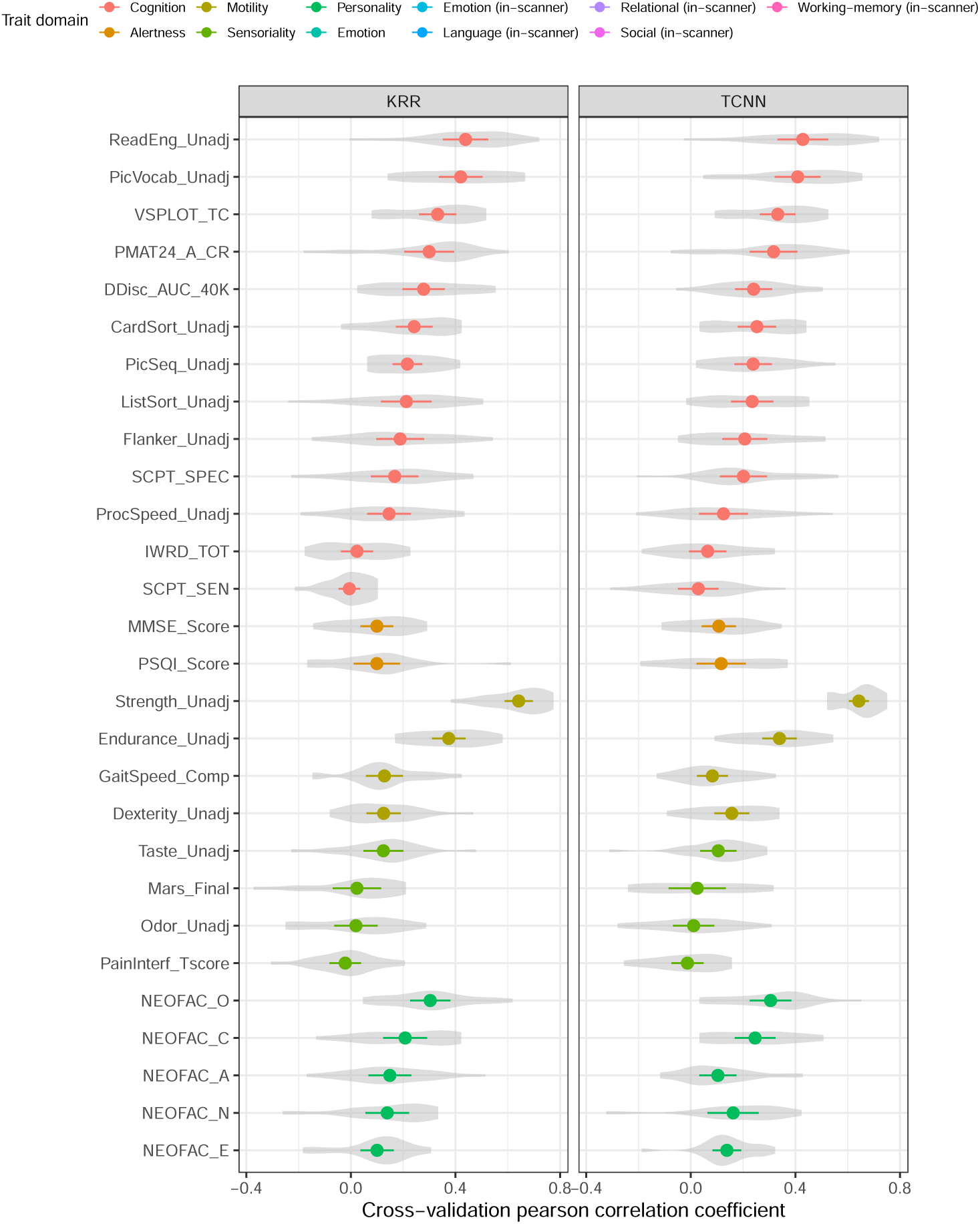
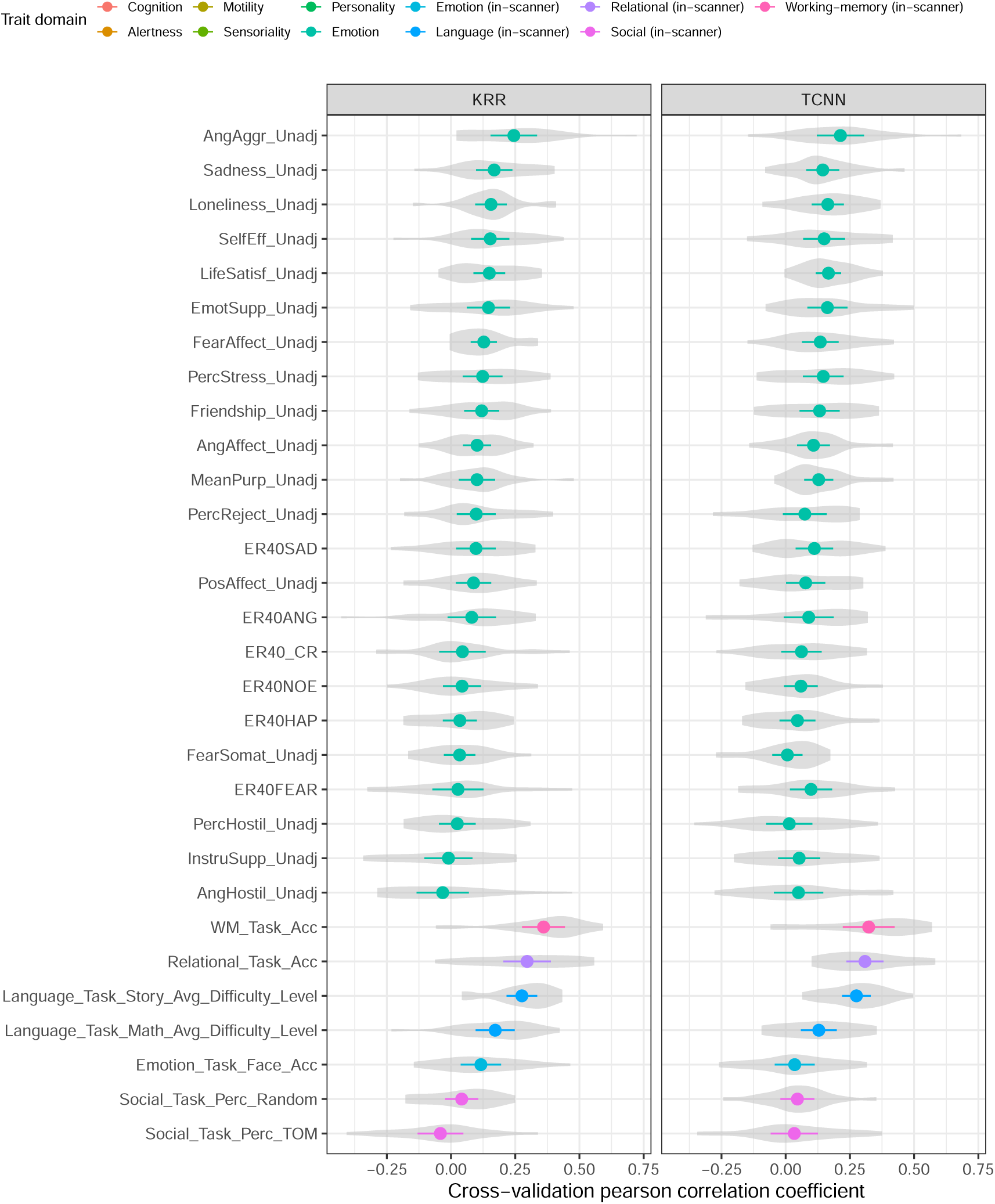
Average performance estimates for 58 behavioral traits in the HCP dataset based on the Pearson correlation coefficient. Confidence intervals are obtained assuming a t-distribution of the mean with 95% confidence, following FCR adjustment.

**Figure S4:**
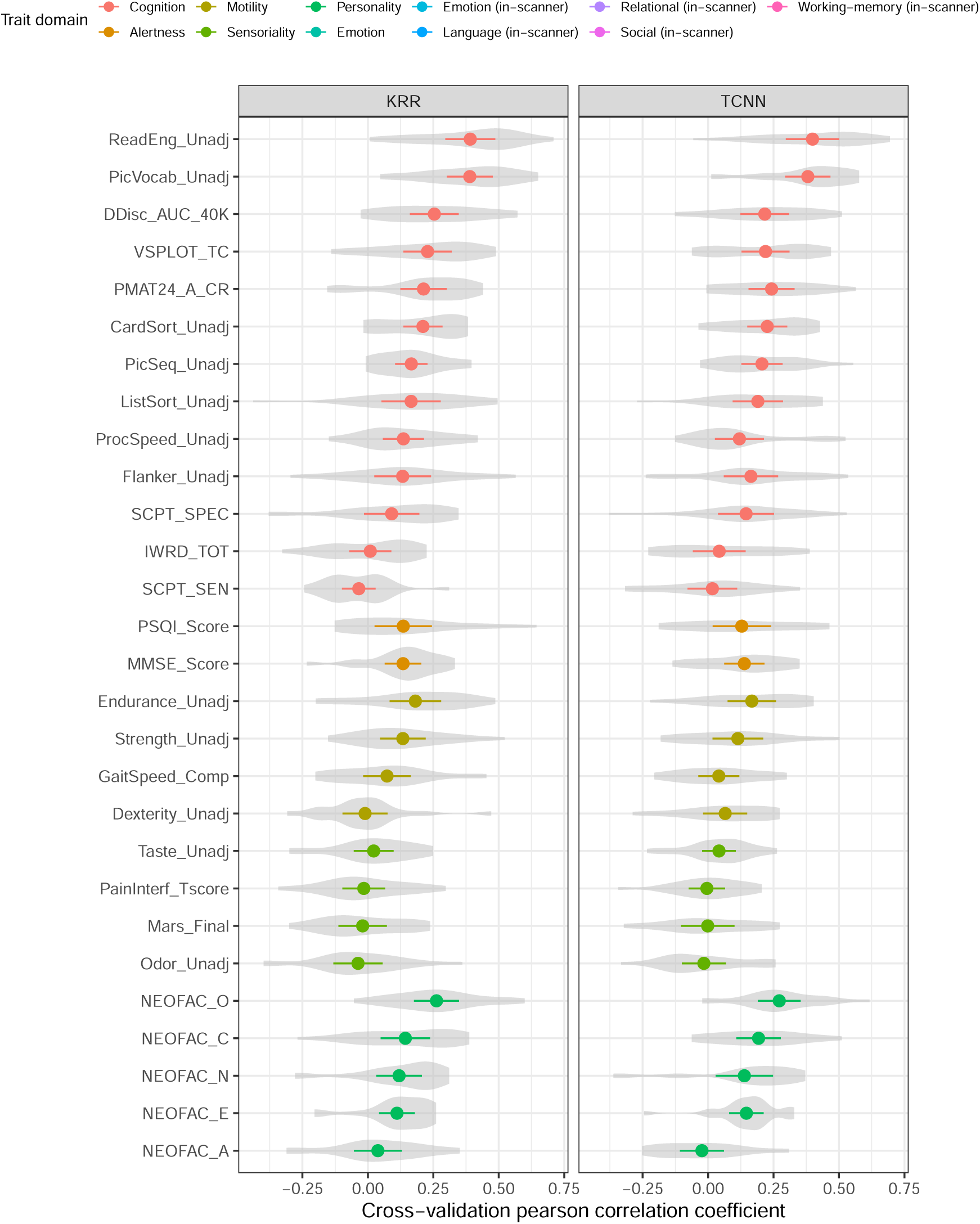
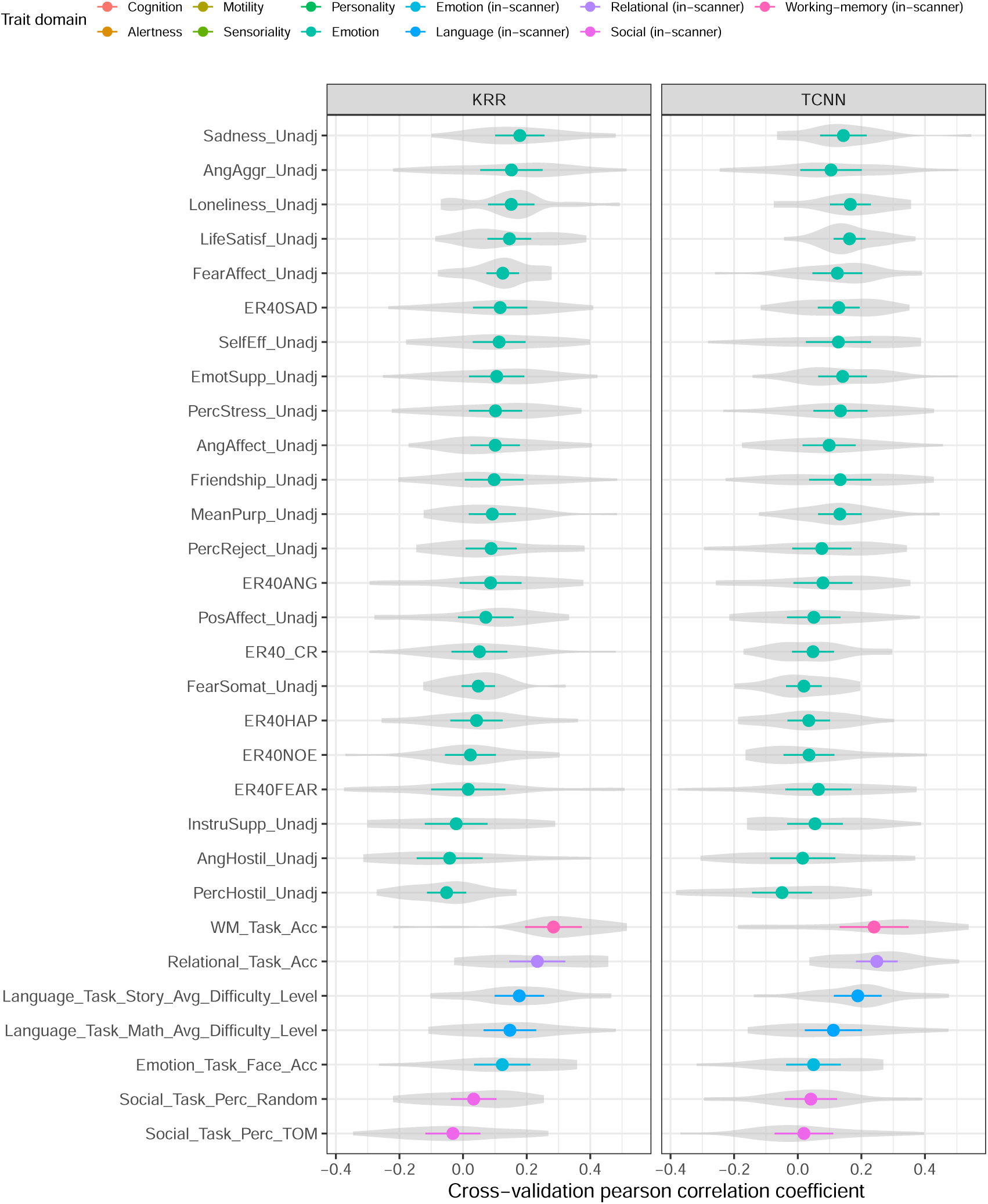
Average performance estimates for 58 behavioral traits in the HCP dataset based on the Pearson correlation coefficient, including covariate regression. Confidence intervals are obtained assuming a t-distribution of the mean with 95% confidence, following FCR adjustment.

**Table S2:**
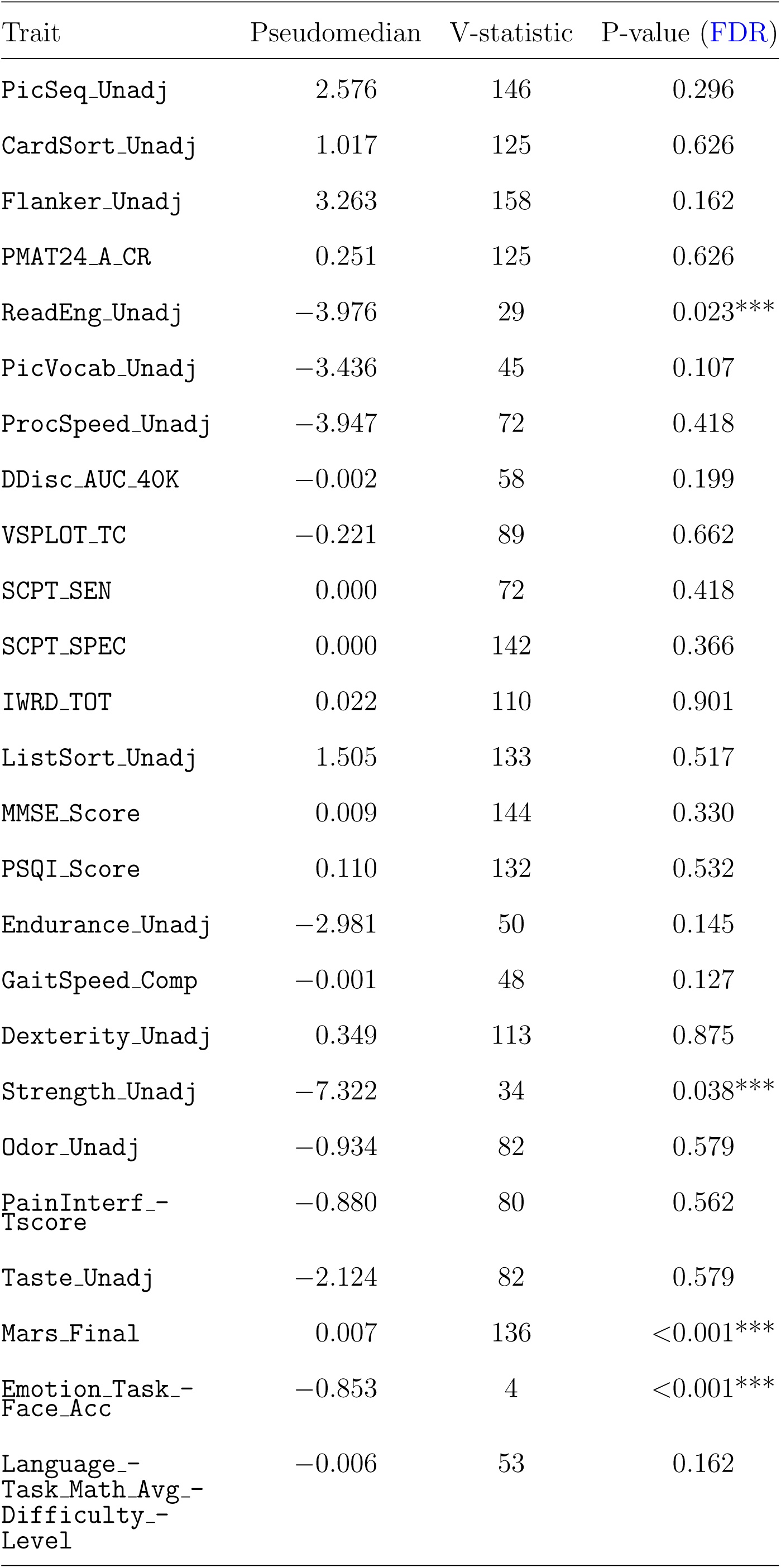
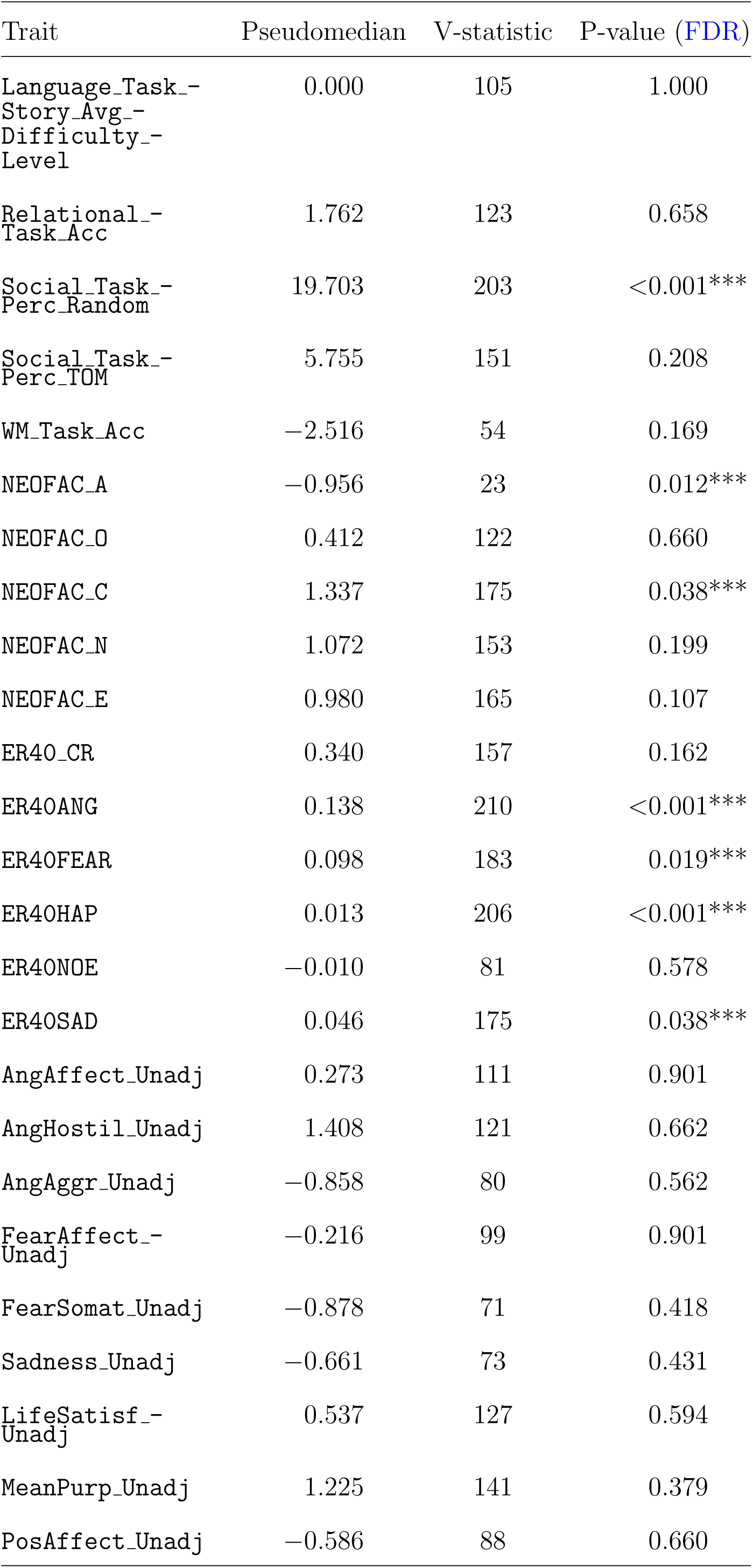
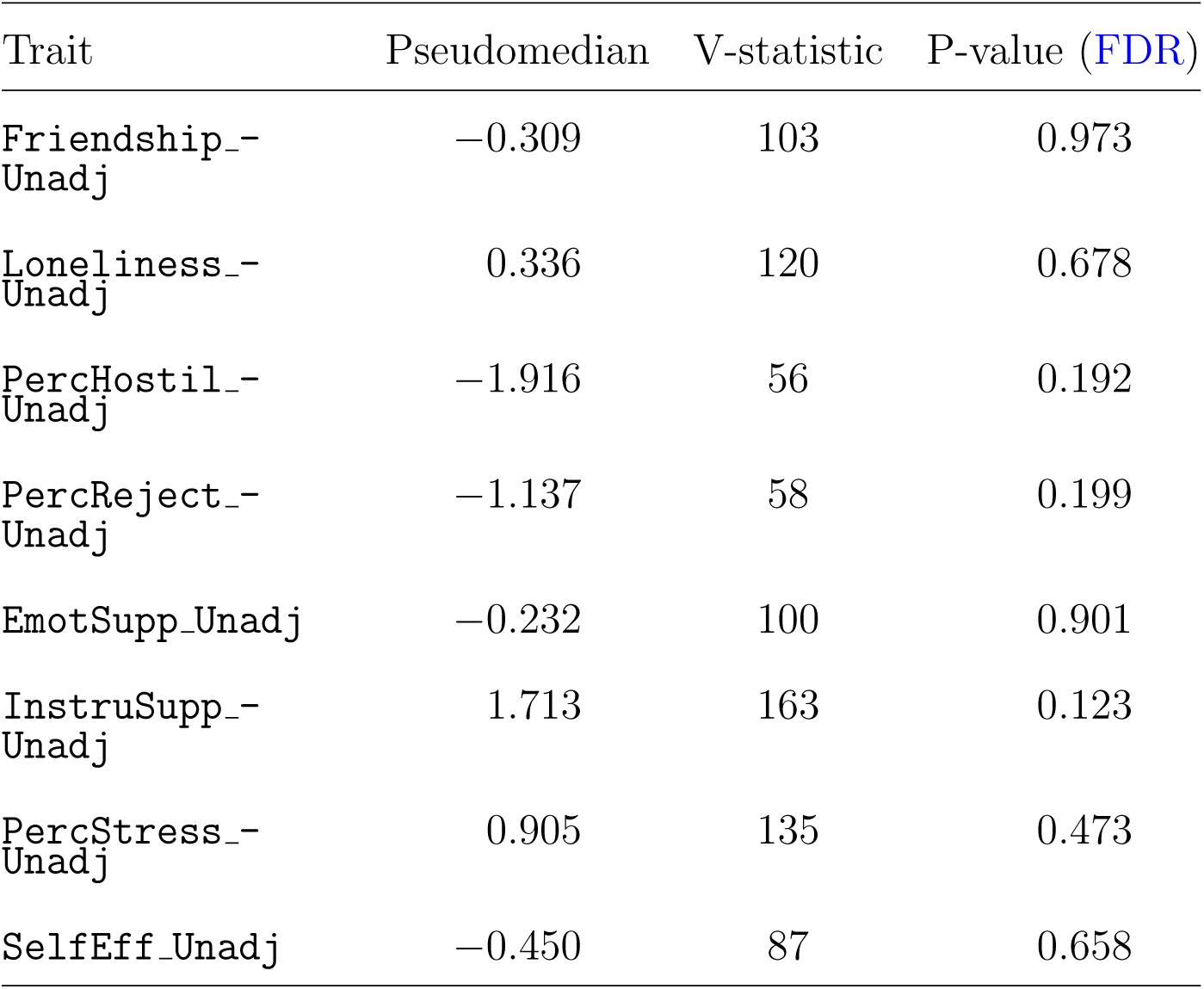
58 behavioral traits in the HCP dataset, without deconfounding. P-values from the pairwise Wilcoxon signed-rank test for the comparison of the TCNN and KRR models in terms of MSE. The V-statistic is the positive rank-sum of the comparison, with a minimum of 0 and a maximum of *N* = (*n*^2^ + *n*)*/*2 = 210 for *n* = 20 (*N* = 136 for Mars Final, with four folds removed). For a comparison between models A and B, if the V-statistic is close to 0, then model A has lower MSE than model B, and if the V-statistic is close to *N*, then model A has higher MSE than model B. A positive pseudomedian indicates that the KRR model has lower MSE than the TCNN model. FCR adjustment was performed.

**Table S3:**
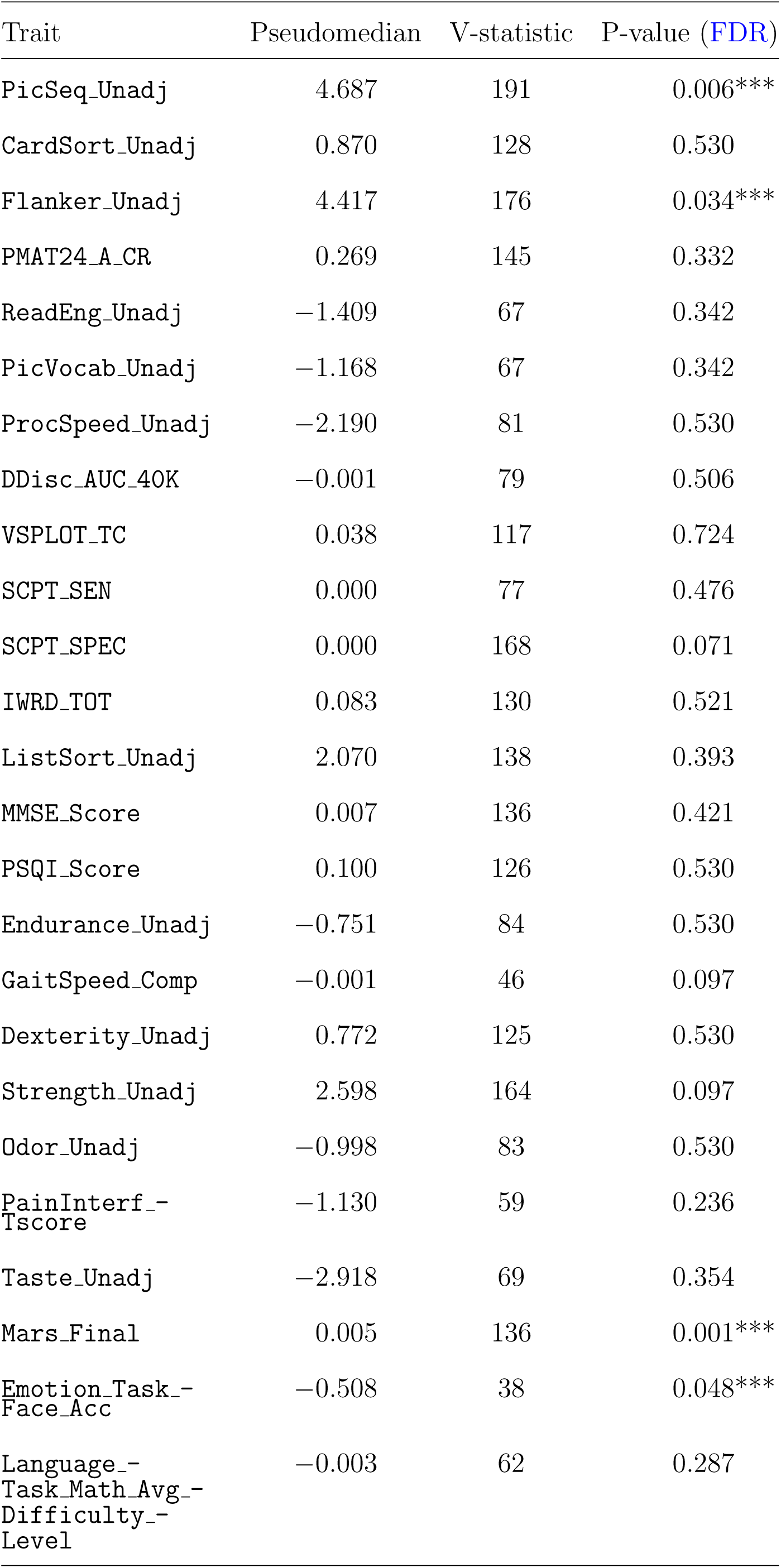
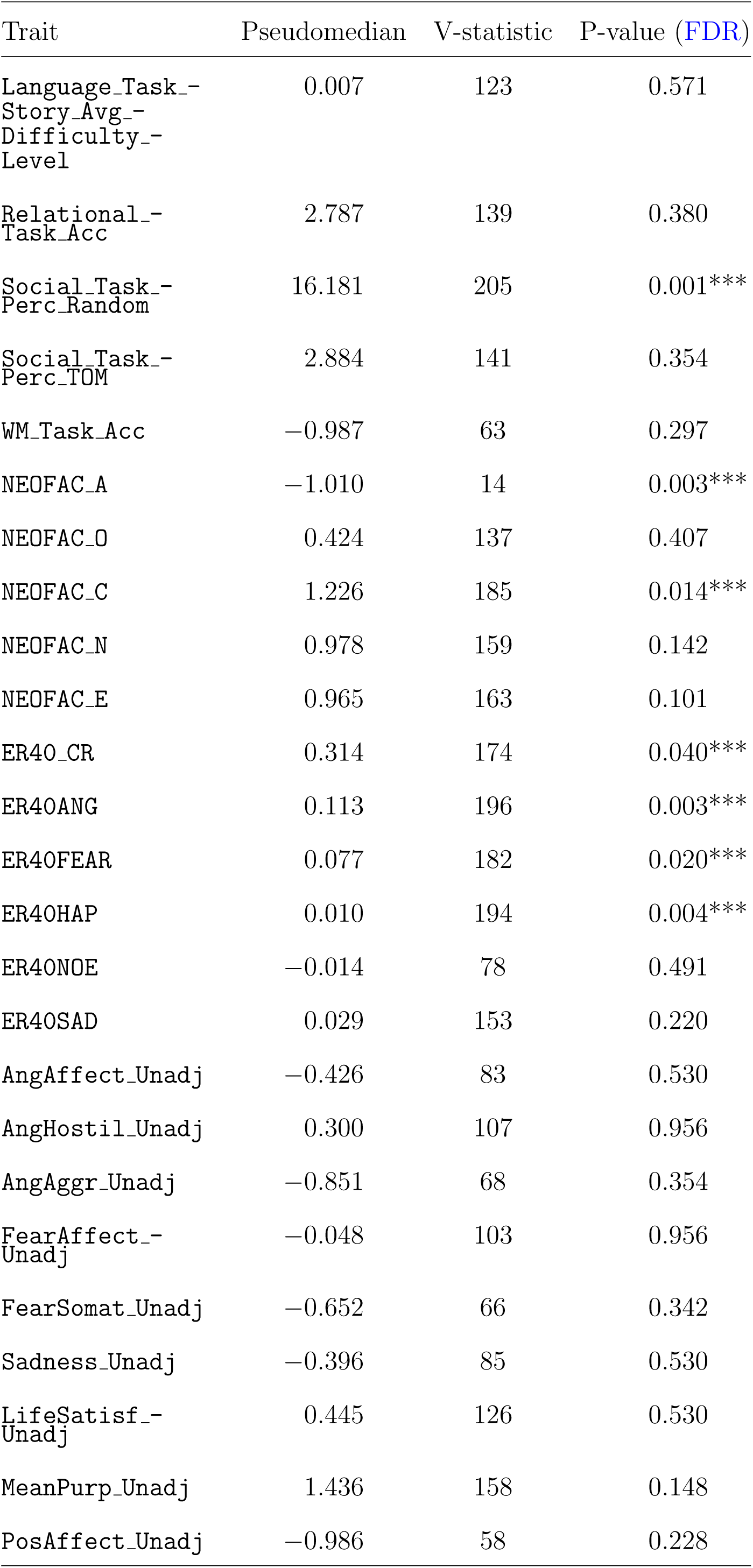
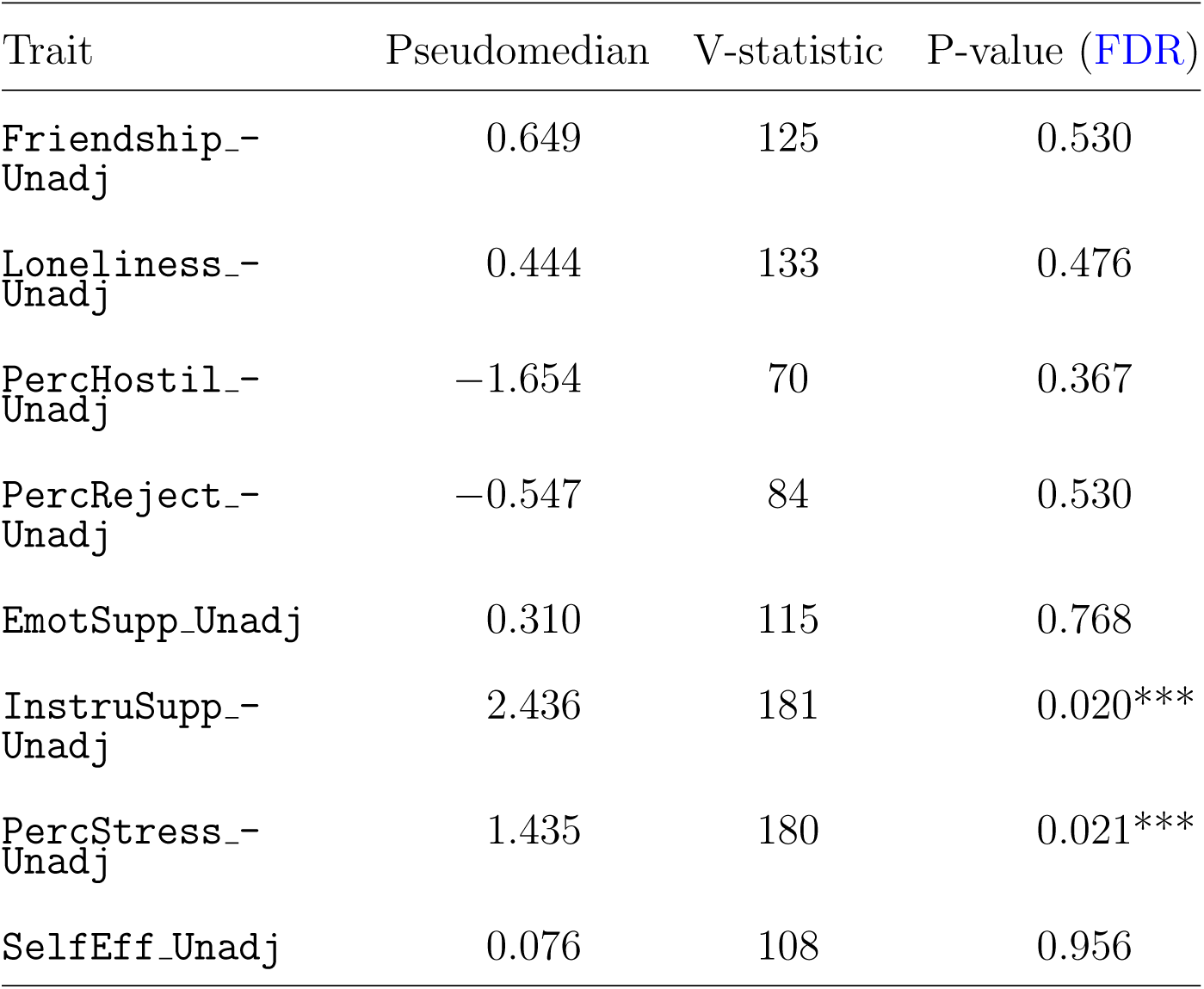
58 behavioral traits in the HCP dataset, including deconfounding. P-values from the pairwise Wilcoxon signed-rank test for the comparison of the TCNN and KRR models in terms of MSE. The V-statistic is the positive rank-sum of the comparison, with a minimum of 0 and a maximum of *N* = (*n*^2^ + *n*)*/*2 = 210 for *n* = 20 (*N* = 136 for Mars Final, with four folds removed). For a comparison between models A and B, if the V-statistic is close to 0, then model A has lower MSE than model B, and if the V-statistic is close to *N*, then model A has higher MSE than model B. A positive pseudomedian indicates that the KRR model has lower MSE than the TCNN model. FCR adjustment was performed.

**Table S4:**
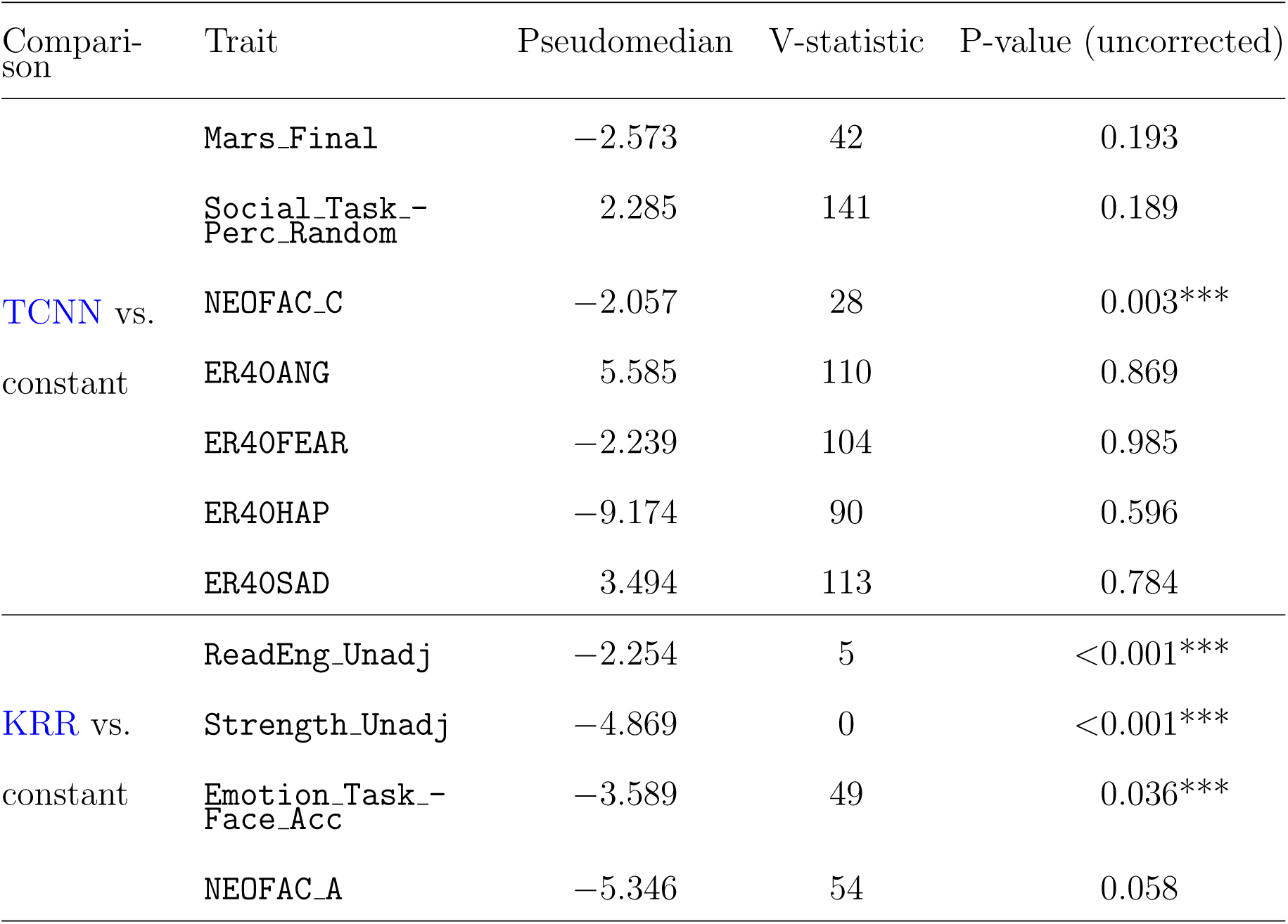
Results from post-hoc pairwise two-sided Wilcoxon signed-rank test for the comparison of the best performing model in Table S2 and the constant model in terms of MSE. The V-statistic is the positive rank-sum of the comparison, with a minimum of 0 and a maximum of *N* = (*n*^2^ + *n*)*/*2 = 210 for *n* = 20 (*N* = 136 for Mars Final, with four folds removed). For a comparison between models A and B, if the V-statistic is close to 0, then model A has lower MSE than model B, and if the V-statistic is close to *N*, then model A has higher MSE than model B. A positive pseudomedian indicates that the constant model has lower MSE than the alternative (TCNN or KRR) model.

**Table S5:** Results from post-hoc pairwise two-sided Wilcoxon signed-rank test for the comparison of the best performing model in Table S3 and the constant model in terms of MSE, including covariate regression. The V-statistic is the positive rank-sum of the comparison, with a minimum of 0 and a maximum of *N* = (*n*^2^ + *n*)*/*2 = 210 for *n* = 20 (*N* = 136 for Mars Final, with four folds removed). For a comparison between models A and B, if the V-statistic is close to 0, then model A has lower MSE than model B, and if the V-statistic is close to *N*, then model A has higher MSE than model B. A positive pseudomedian indicates that the constant model has lower MSE than the alternative (TCNN or KRR) model.

| Comparison | Trait | Pseudomedian | V-statistic | P-value (uncorrected) |
| --- | --- | --- | --- | --- |
| TCNN vs.<br>constant | PicSeq_Unadj | −42.773 | 0 | <0.001*** |
|  | Flanker_Unadj | −24.295 | 0 | <0.001*** |
|  | Mars_Final | −0.002 | 0 | <0.001*** |
|  | Social_Task_-<br>Perc_Random | −14.361 | 2 | <0.001*** |
|  | NEOFAC_C | −6.895 | 0 | <0.001*** |
|  | ER40_CR | −0.838 | 1 | <0.001*** |
|  | ER40ANG | −0.181 | 1 | <0.001*** |
|  | ER40FEAR | −0.188 | 2 | <0.001*** |
|  | ER40HAP | −0.008 | 0 | <0.001*** |
|  | InstruSupp_-<br>Unadj | −10.350 | 2 | <0.001*** |
|  | PercStress_-<br>Unadj | −13.317 | 0 | <0.001*** |
| KRR vs.<br>constant | Emotion_Task_-<br>Face_Acc | −1.287 | 74 | 0.261 |
|  | NEOFAC_A | −6.425 | 0 | <0.001*** |

## E Scalability of models

Figures S5 and S6 depict the fold-wise paired difference in MSE between the TCNN and KRR models, as functions of the number of subjects in the training set and the number of time points available in the validation set. For each figure, a regression of the form *y* = *β*_0_ +*β*_diff_ *·* log(*x*) was estimated, where *x* is the number of subjects or time points, and *y* is the performance difference. Non-significant traits are ommitted from the plots, and since no differences were found in the Pearson’s correlation coefficient, these figures are not shown for this measure. Figures S7 and S8 show the performance of the TCNN and KRR models in terms of the Pearson’s correlation coefficient between predicted and true values, as a function of the number of subjects in the training set and the number of time points available in the validation set, respectively. Figures S9 and S10 show the same results in terms of the MSE between predicted and true values. Results are shown for the raw data, without covariate regression.

**Figure S5:**
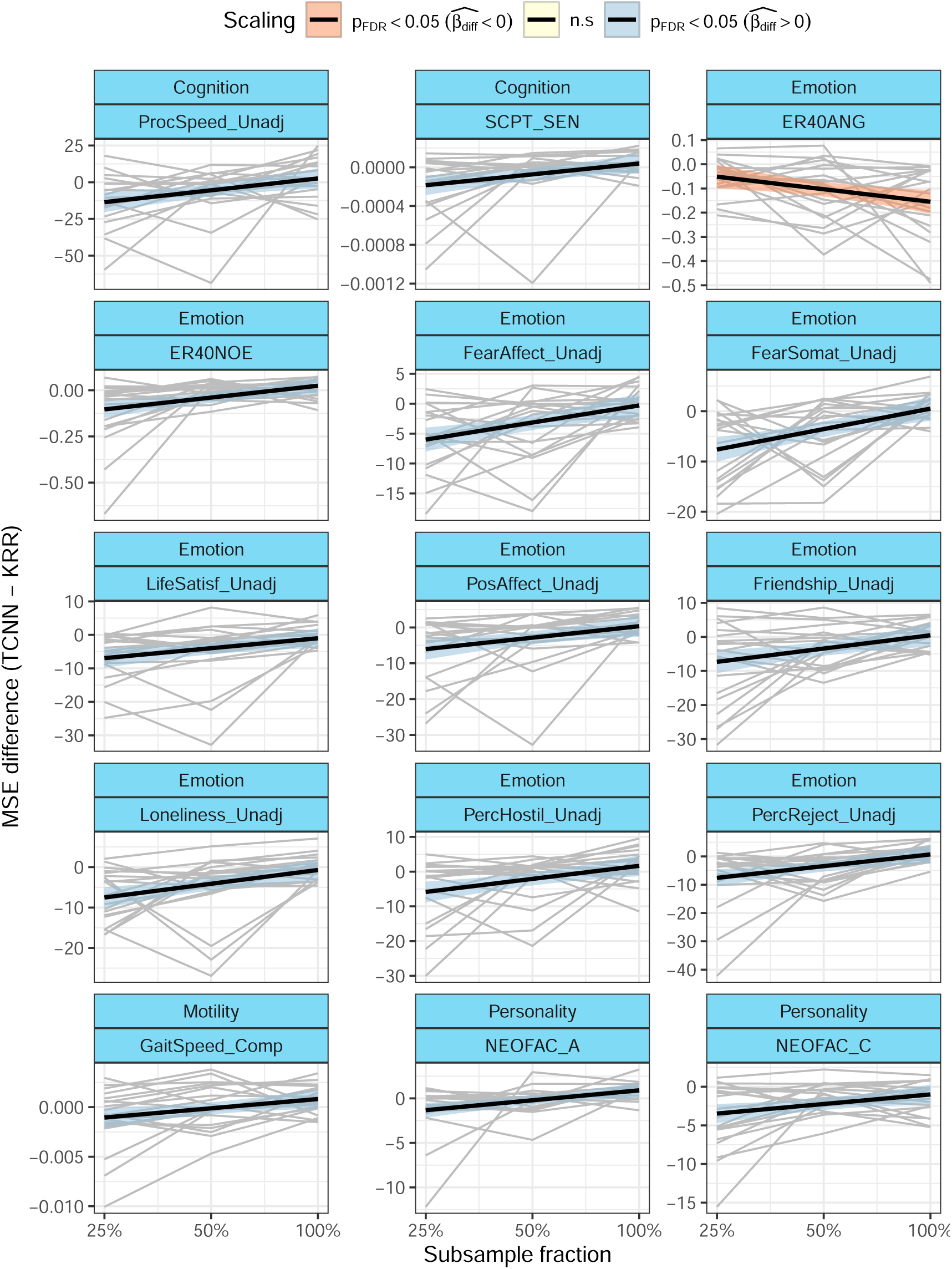
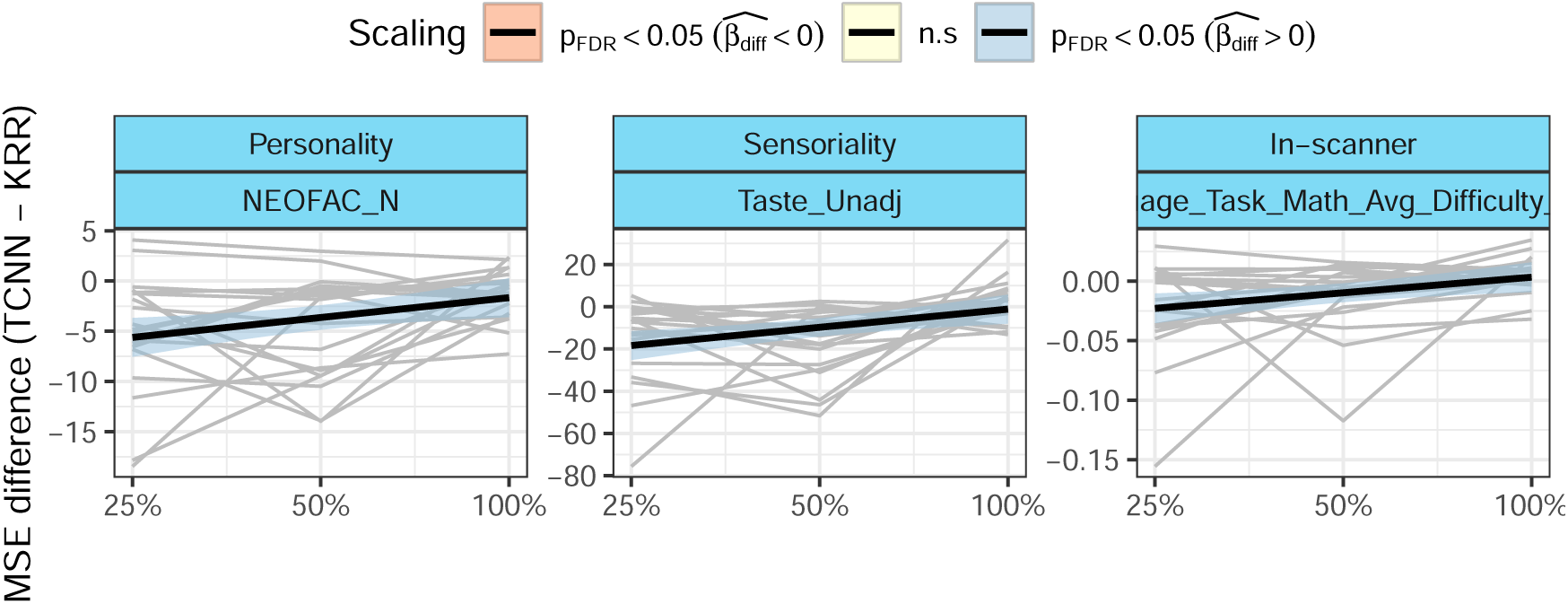
Scaling of the MSE difference with the number of subjects in the training sets.

**Figure S6:**
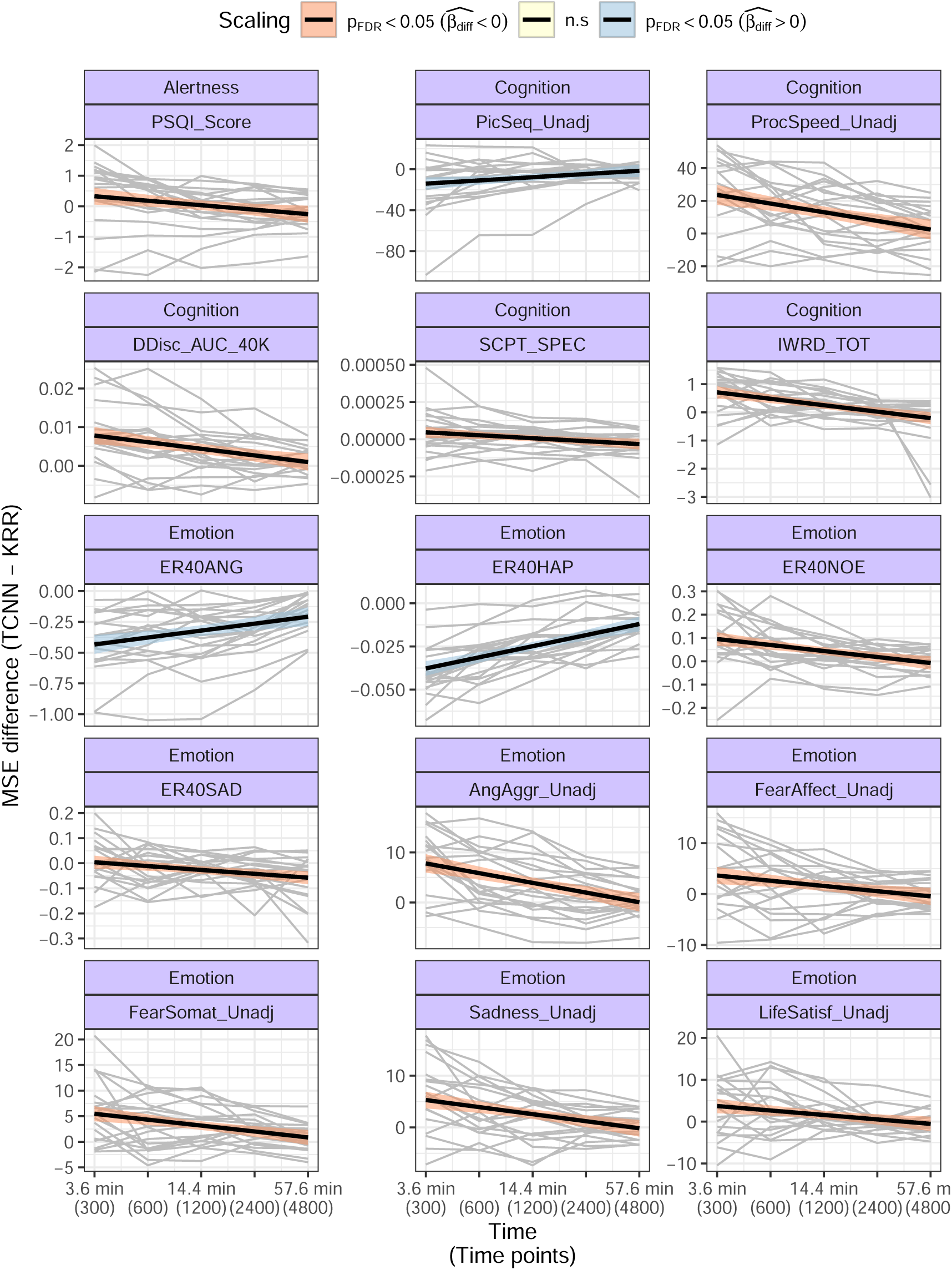
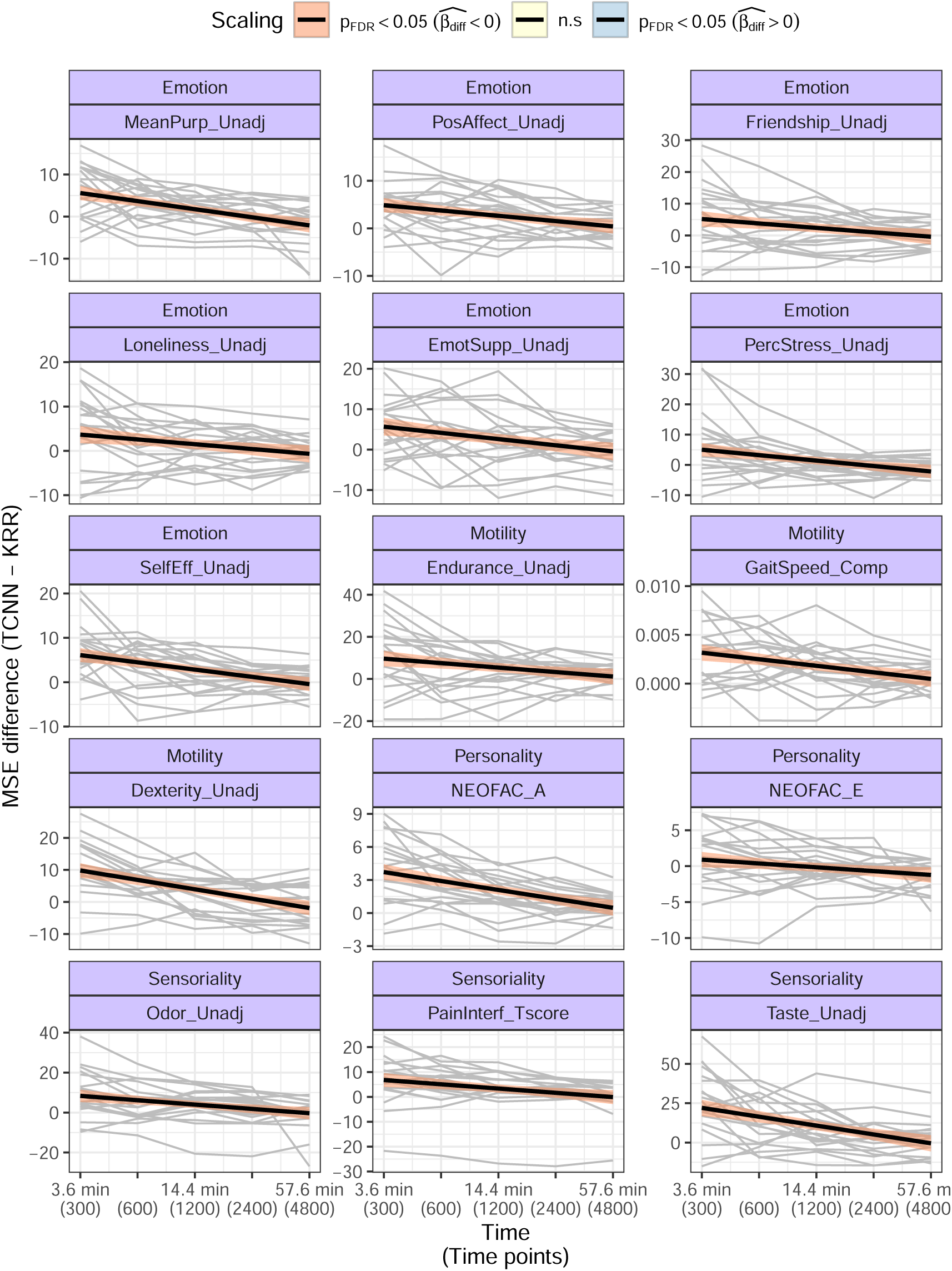
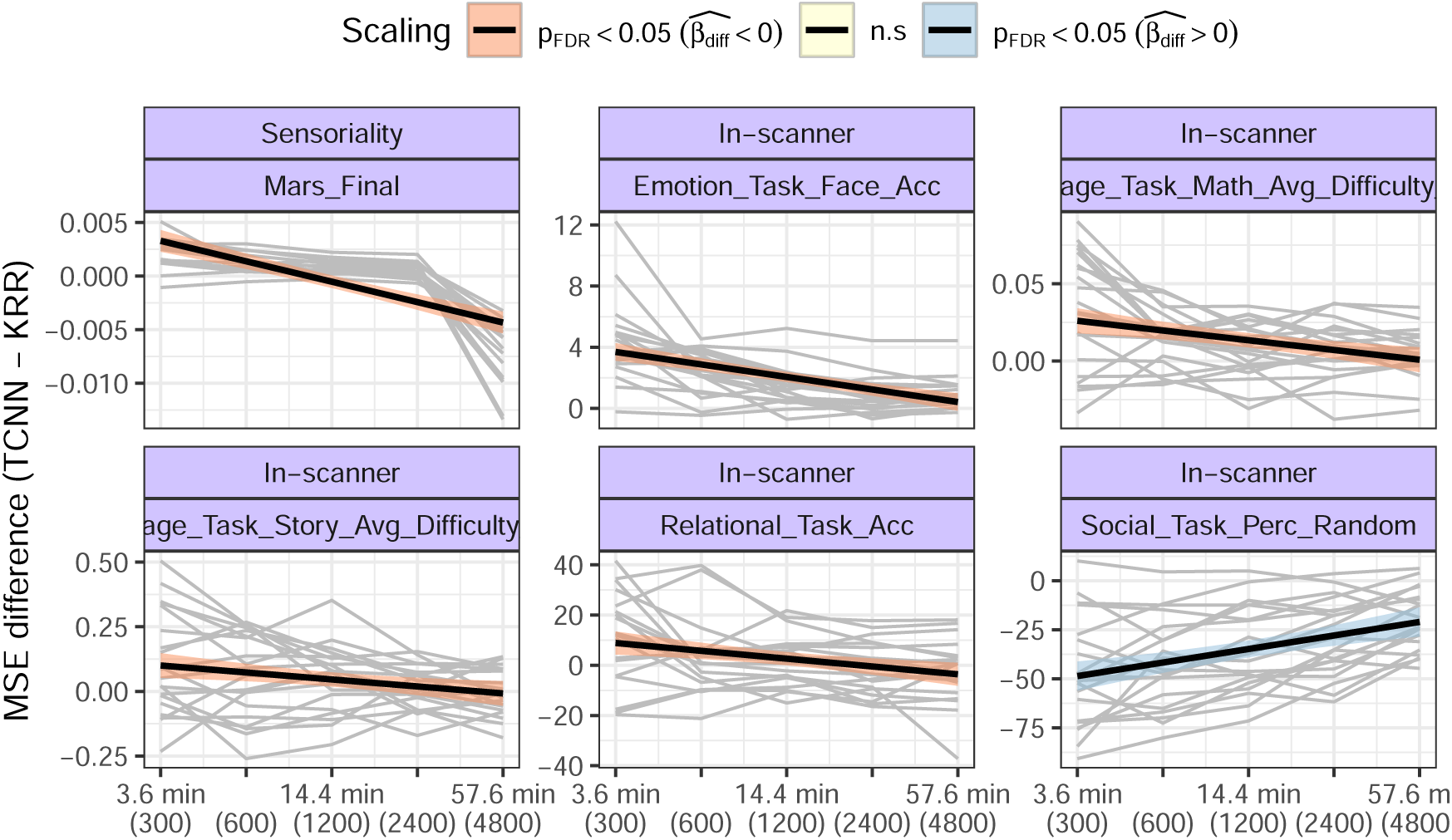
Scaling of the MSE difference with the length of the time series in the validation sets.

**Figure S7:**
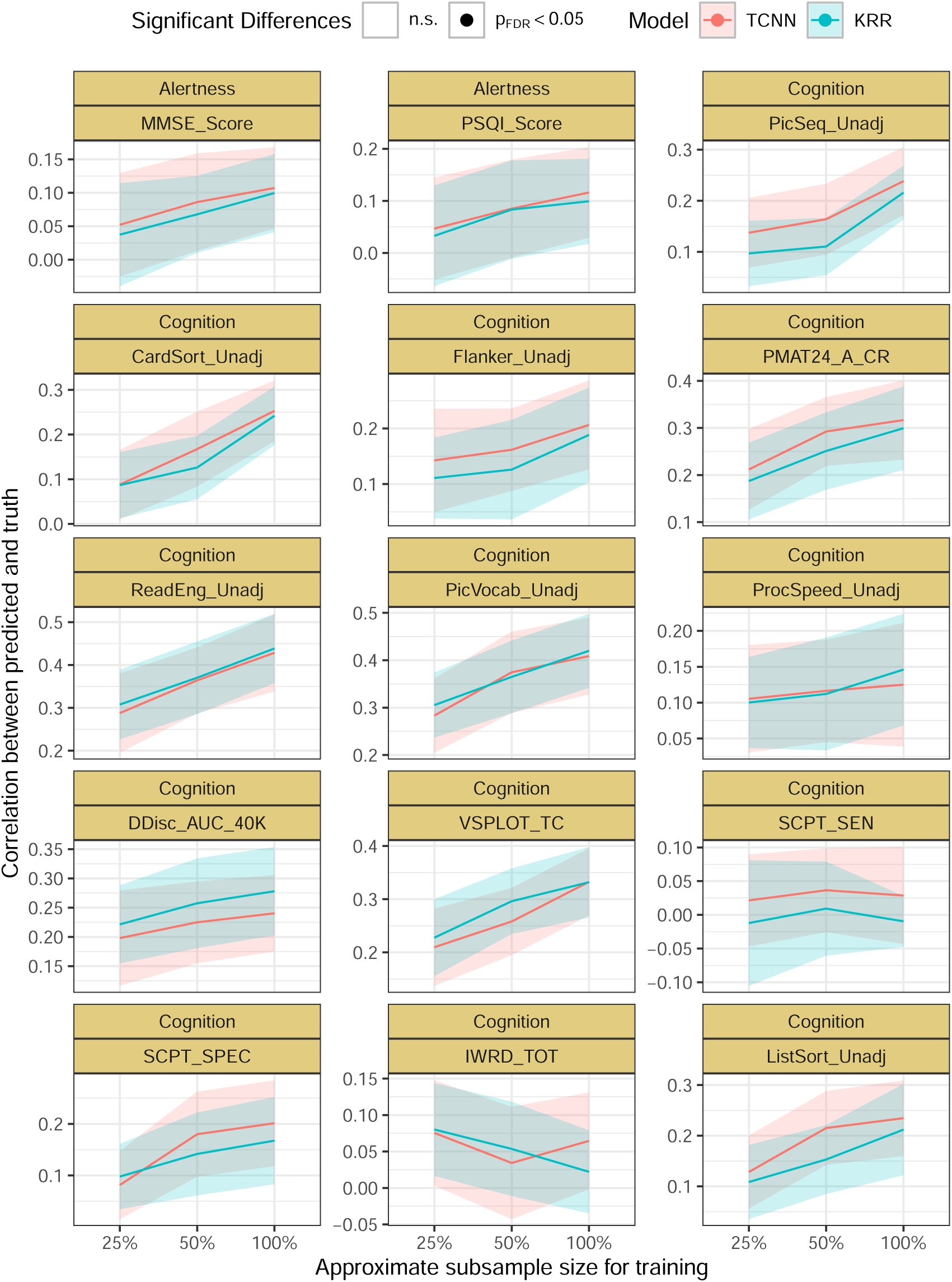
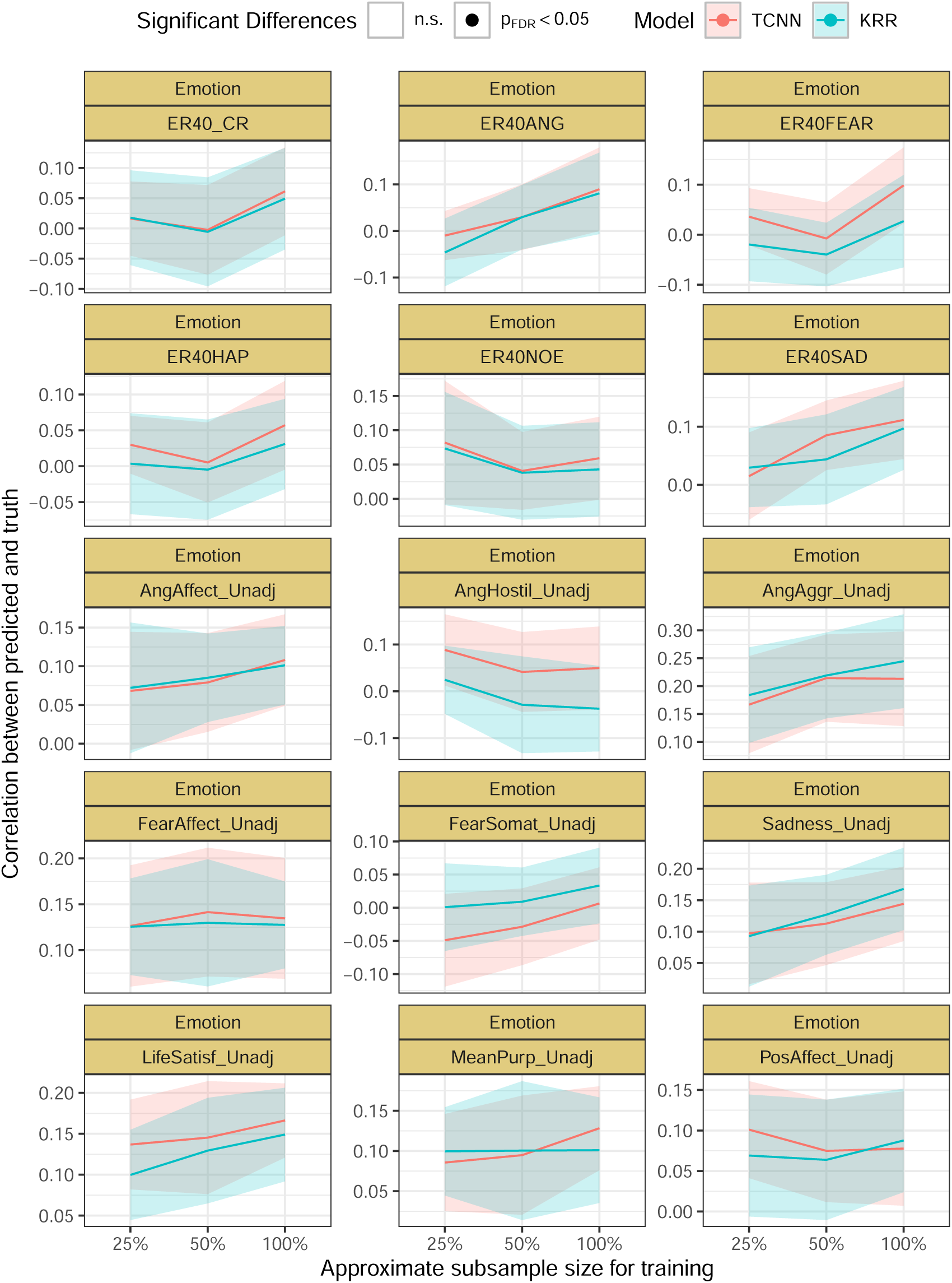
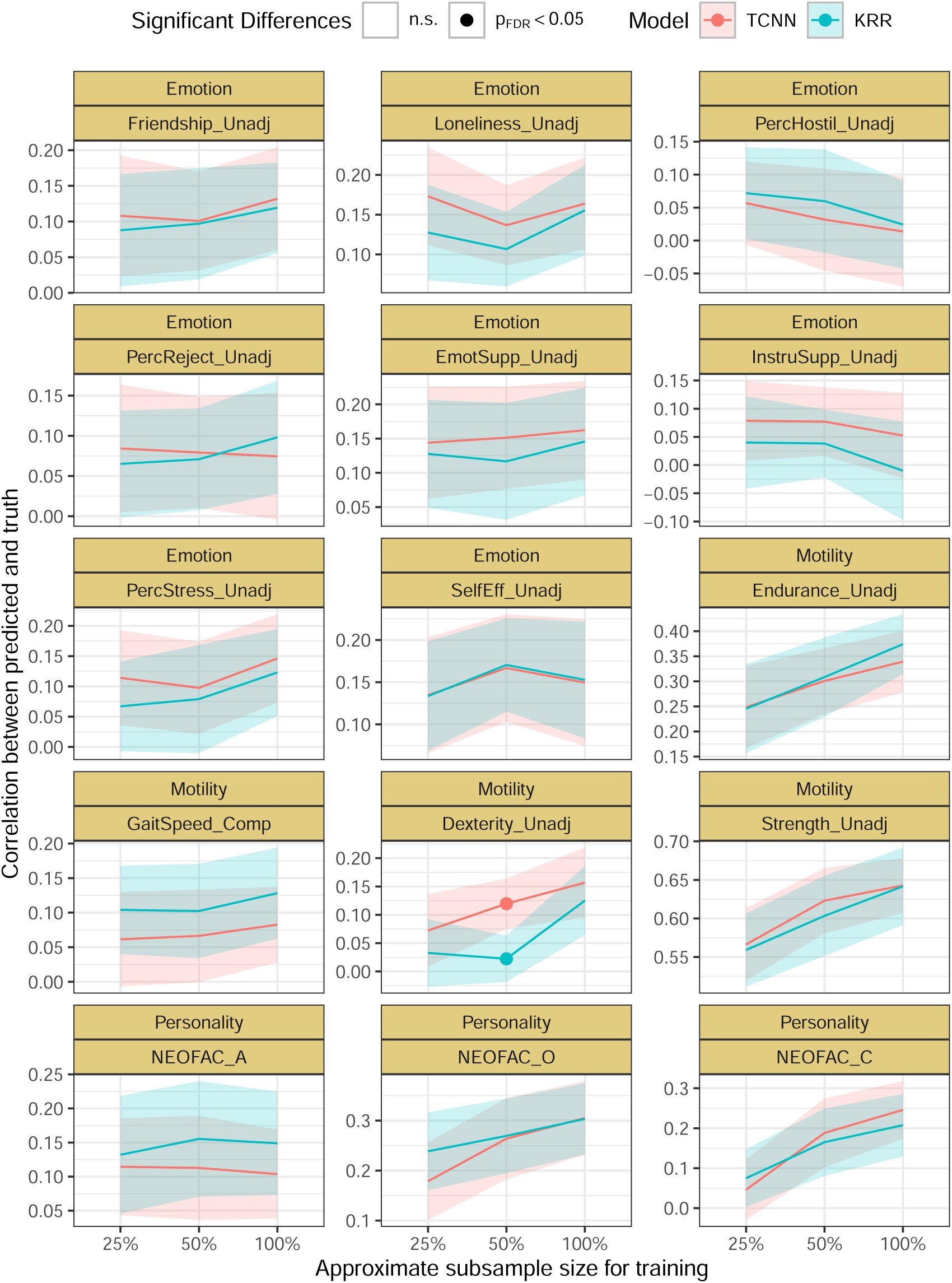
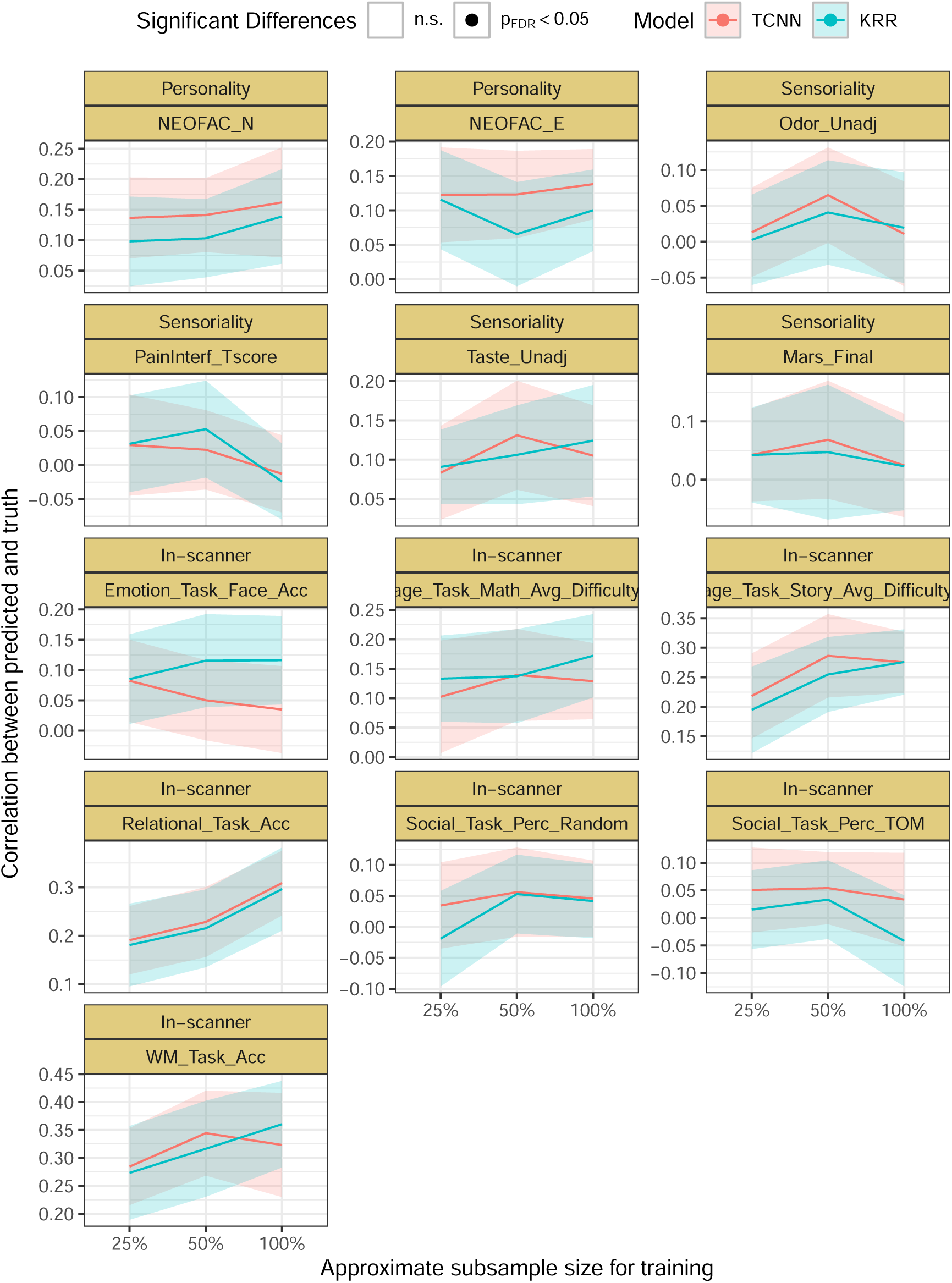
Scaling of the Pearson’s correlation coefficient between predicted and true values with the number of subjects in the training sets.

**Figure S8:**
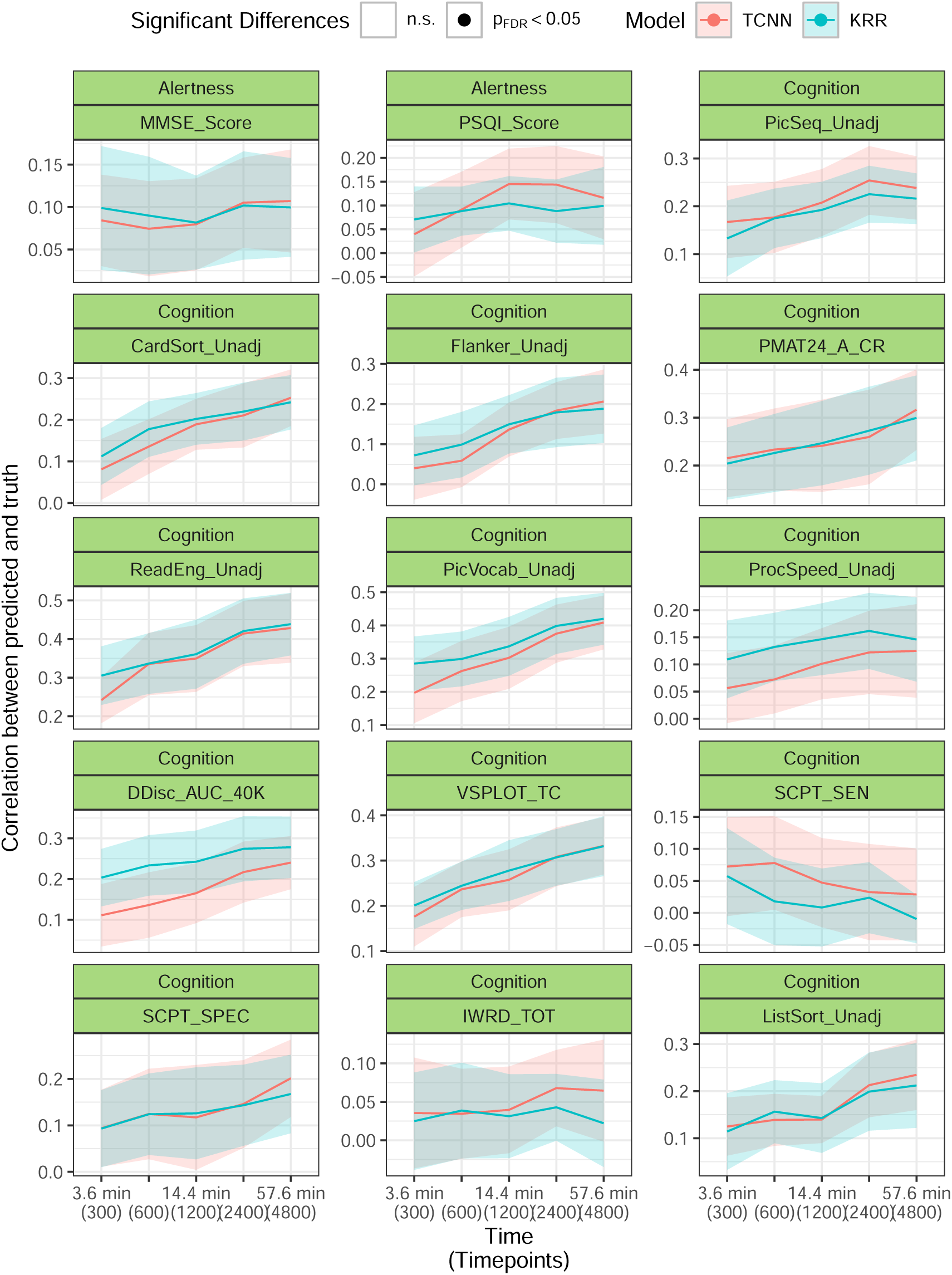
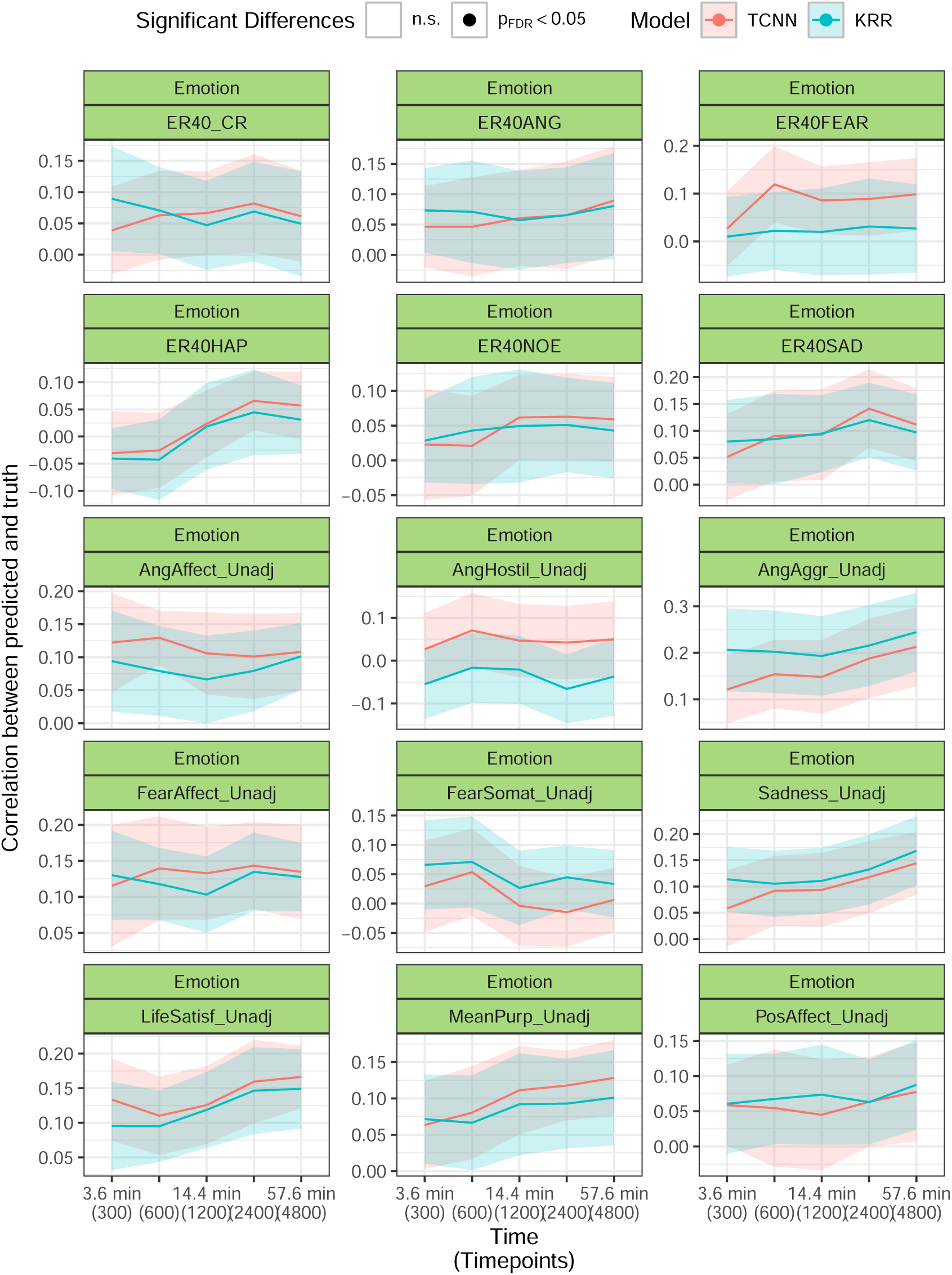
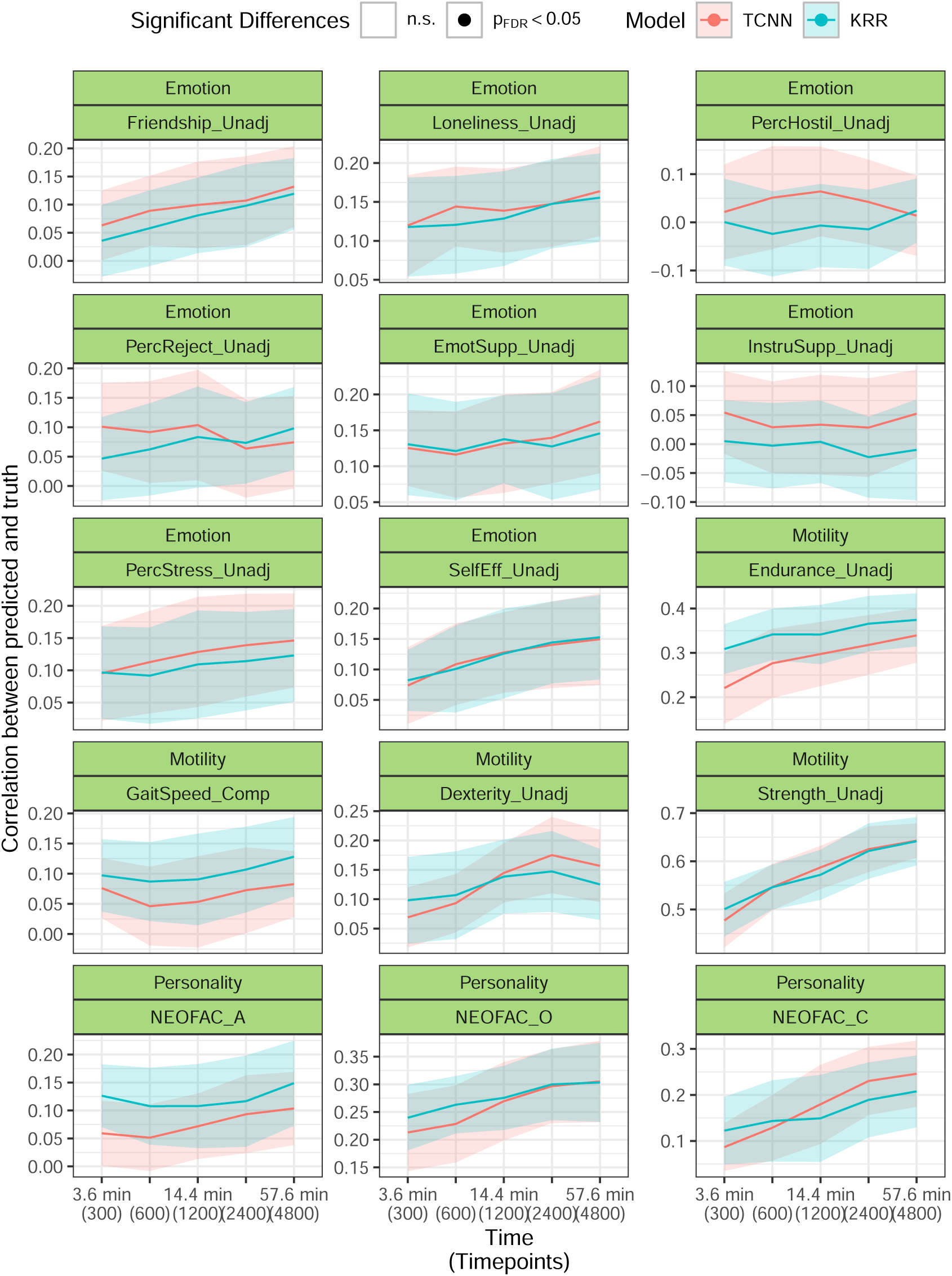
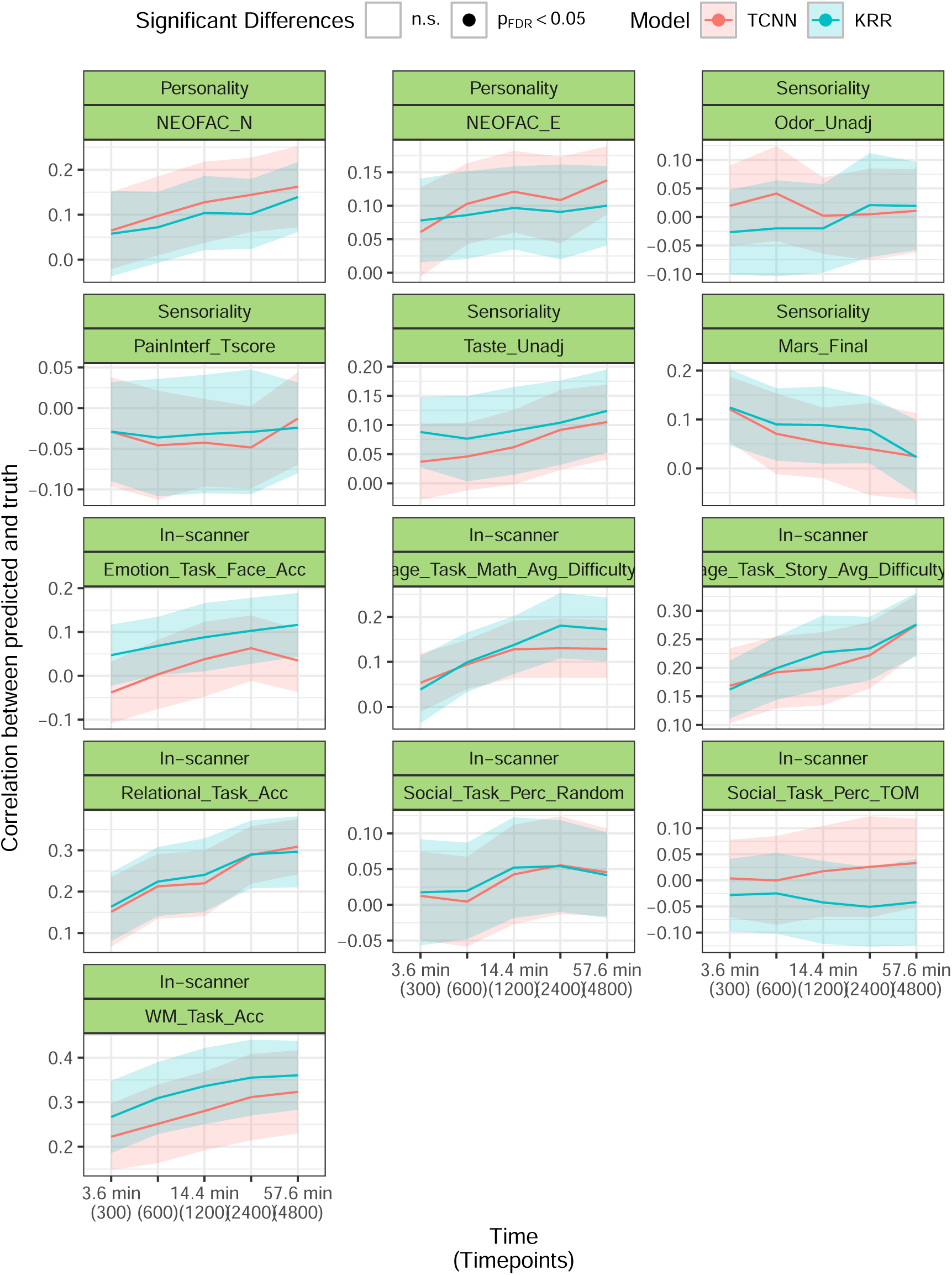
Scaling of the Pearson’s correlation coefficient between predicted and true values with the length of the time series in the validation sets.

**Figure S9:**
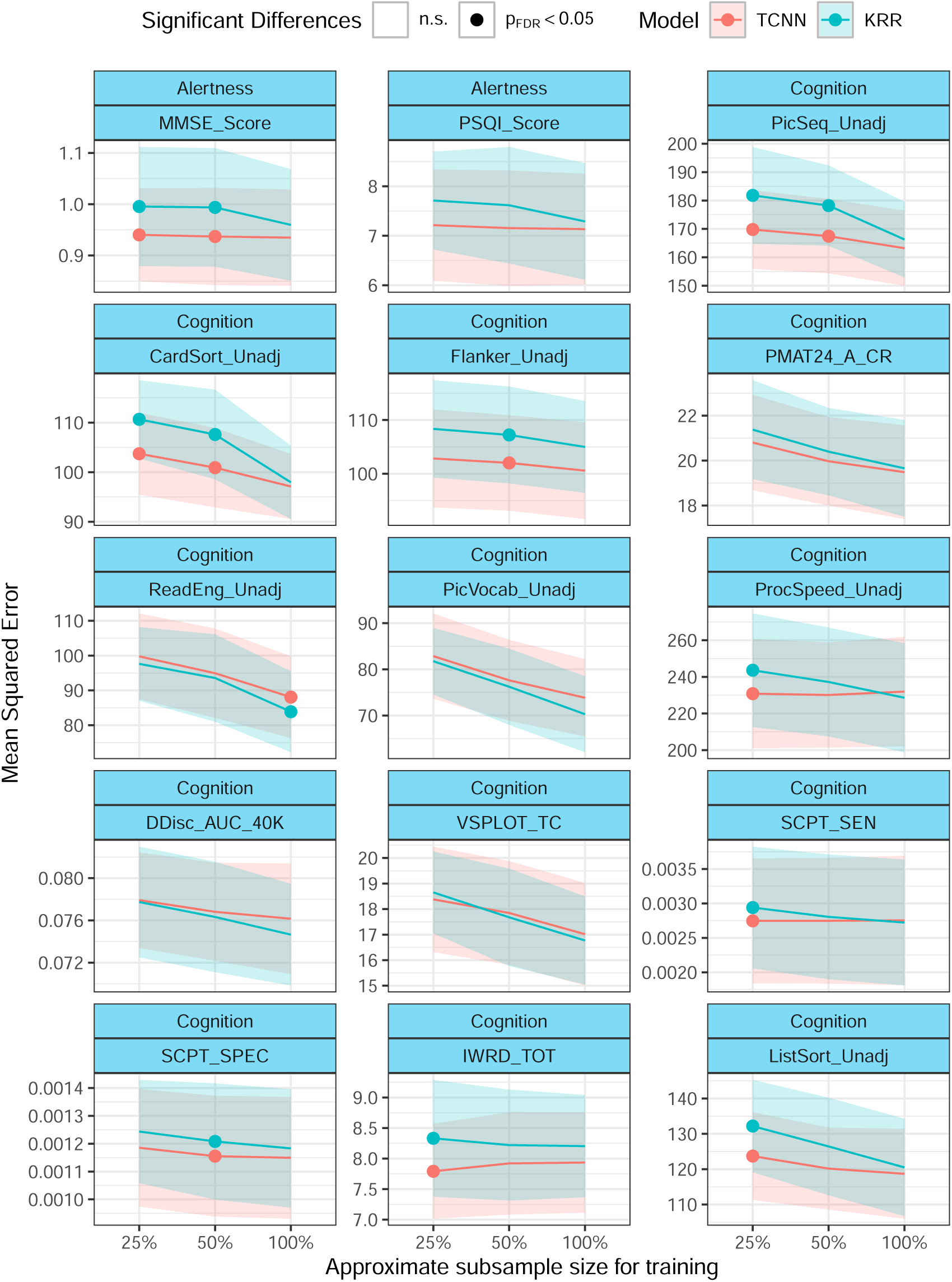
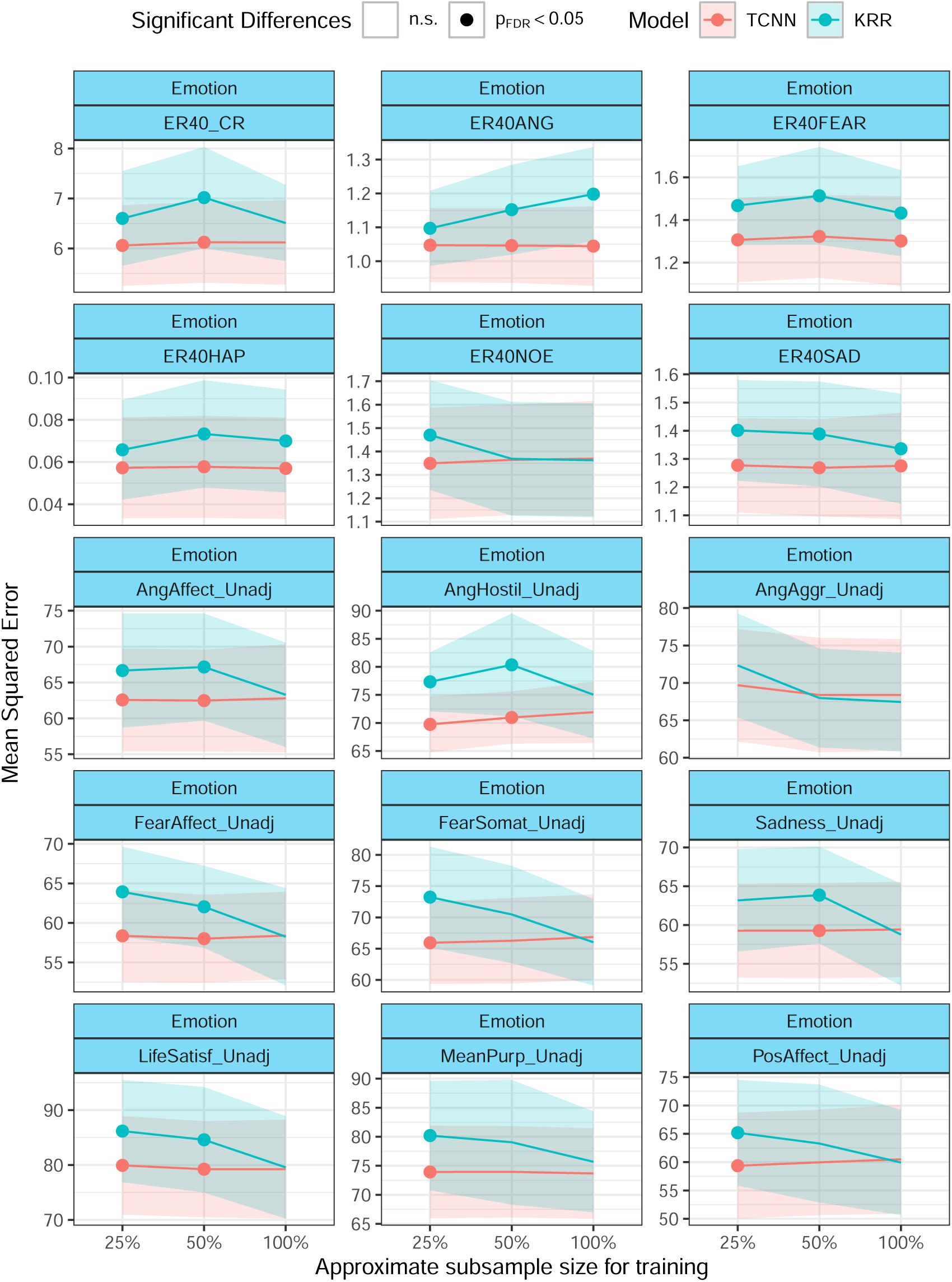
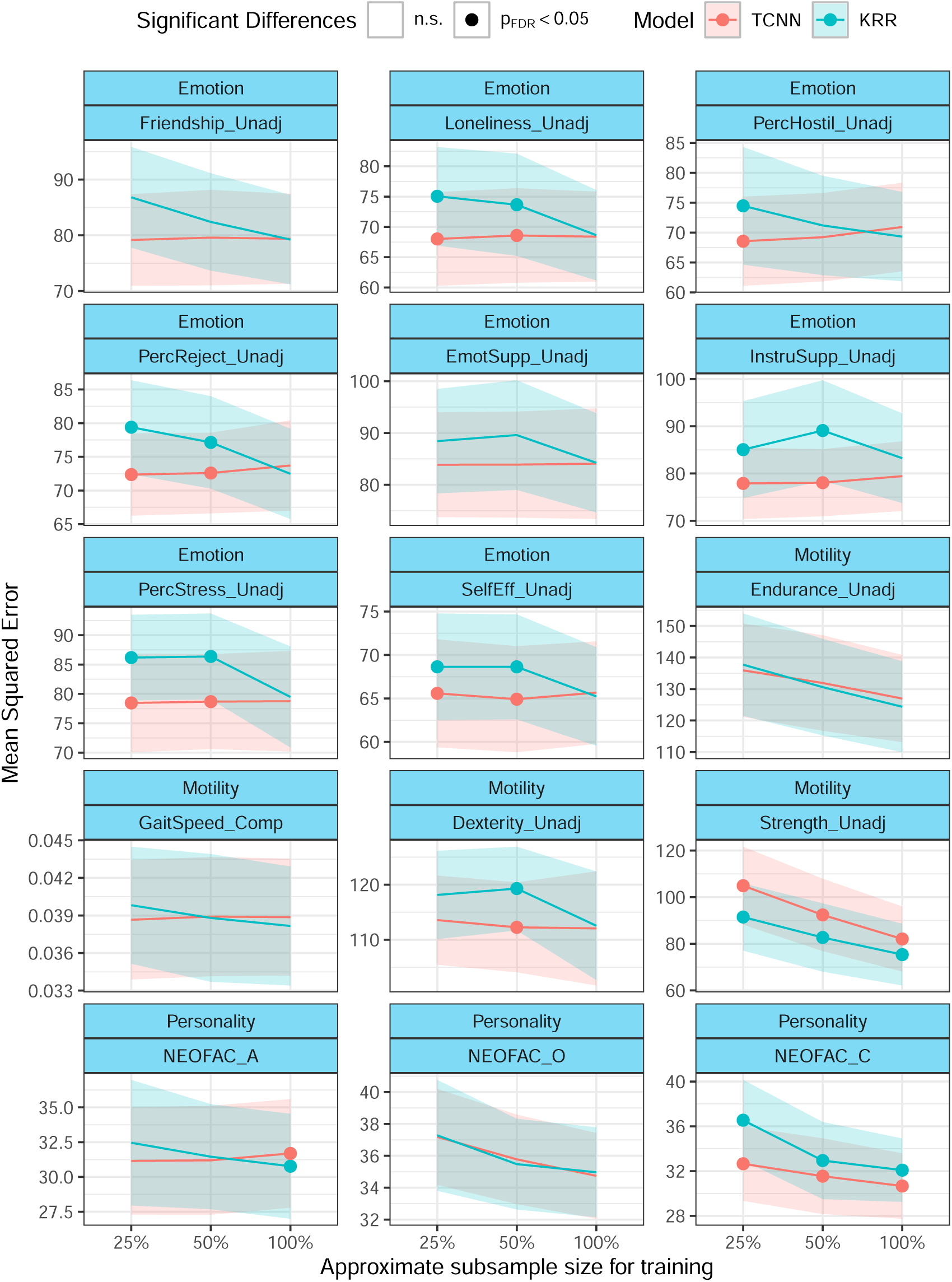
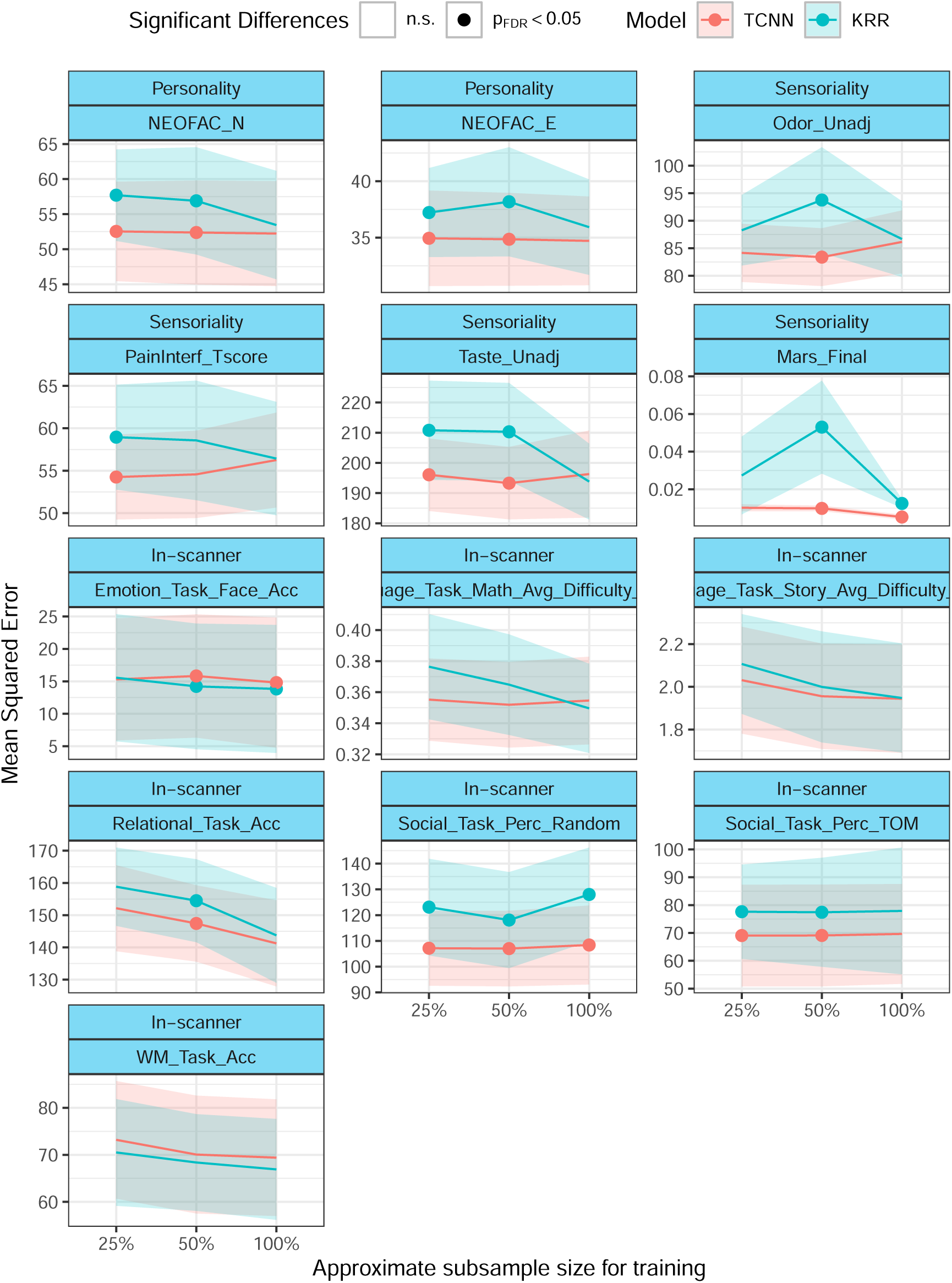
Scaling of the MSE between predicted and true values with the number of subjects in the training sets.

**Figure S10:**
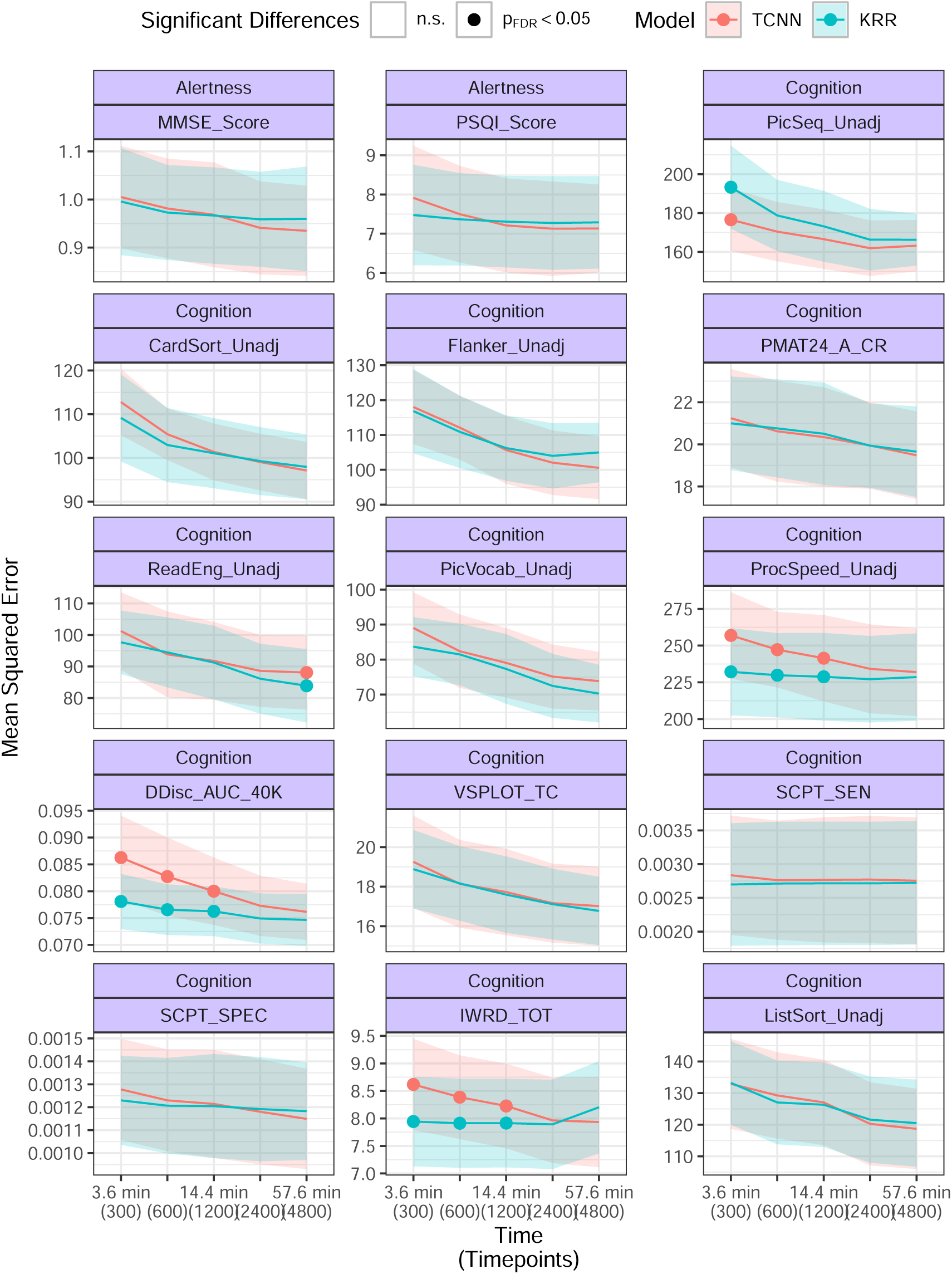
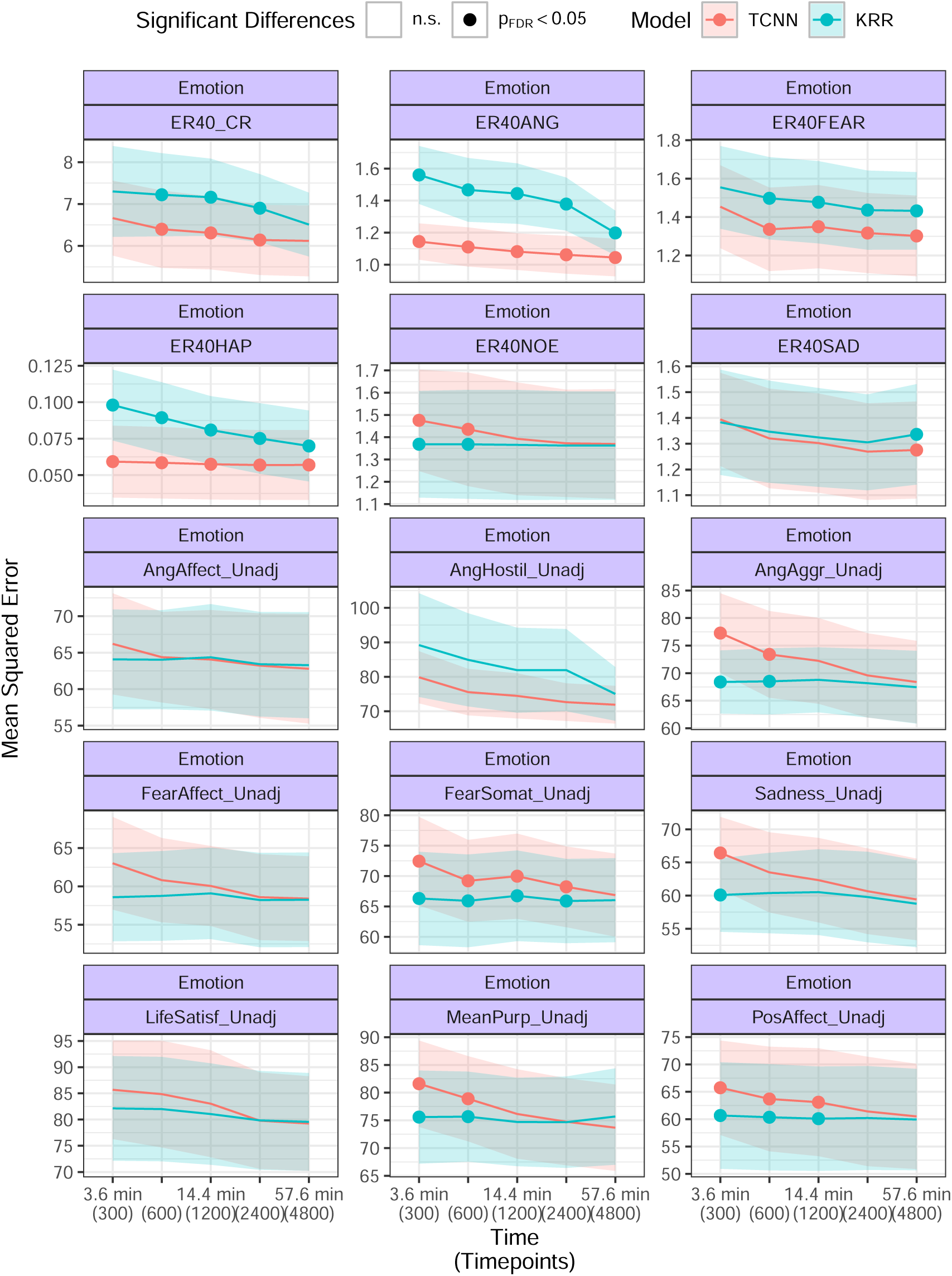
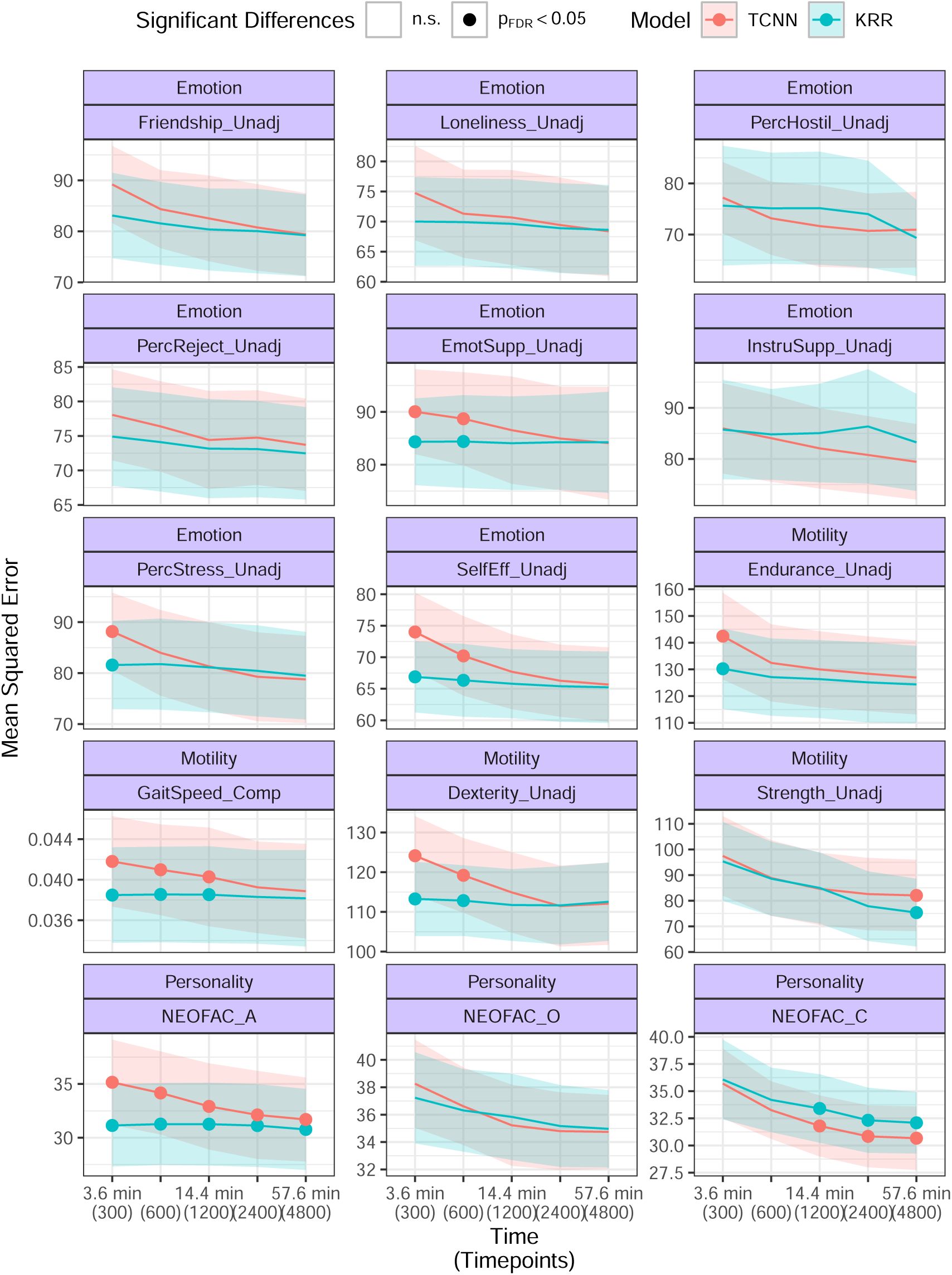
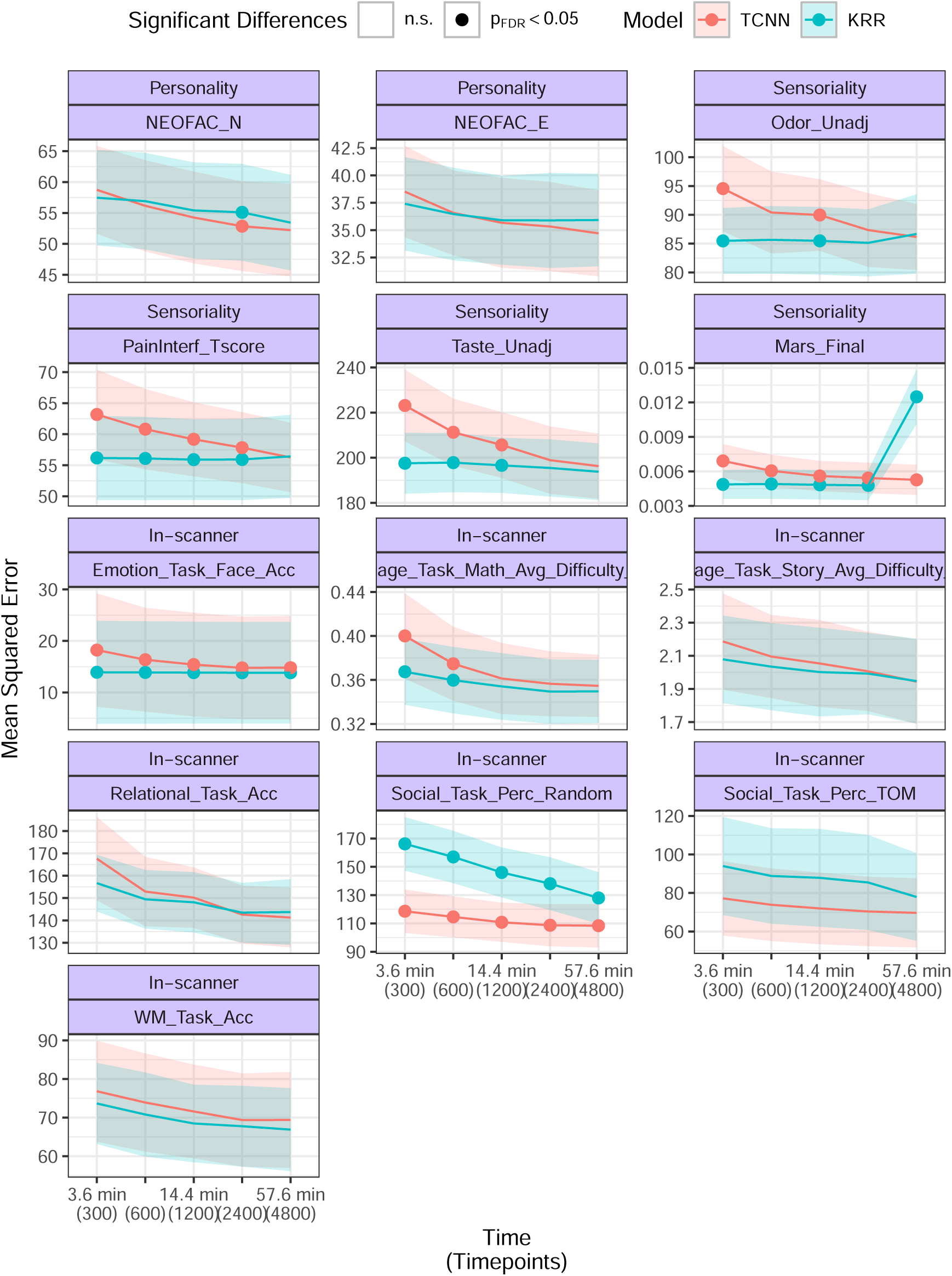
Scaling of the MSE between predicted and true values with the length of the time series in the validation sets.

## F Effect of covariate regression on performance estimates

**Figure S11:**
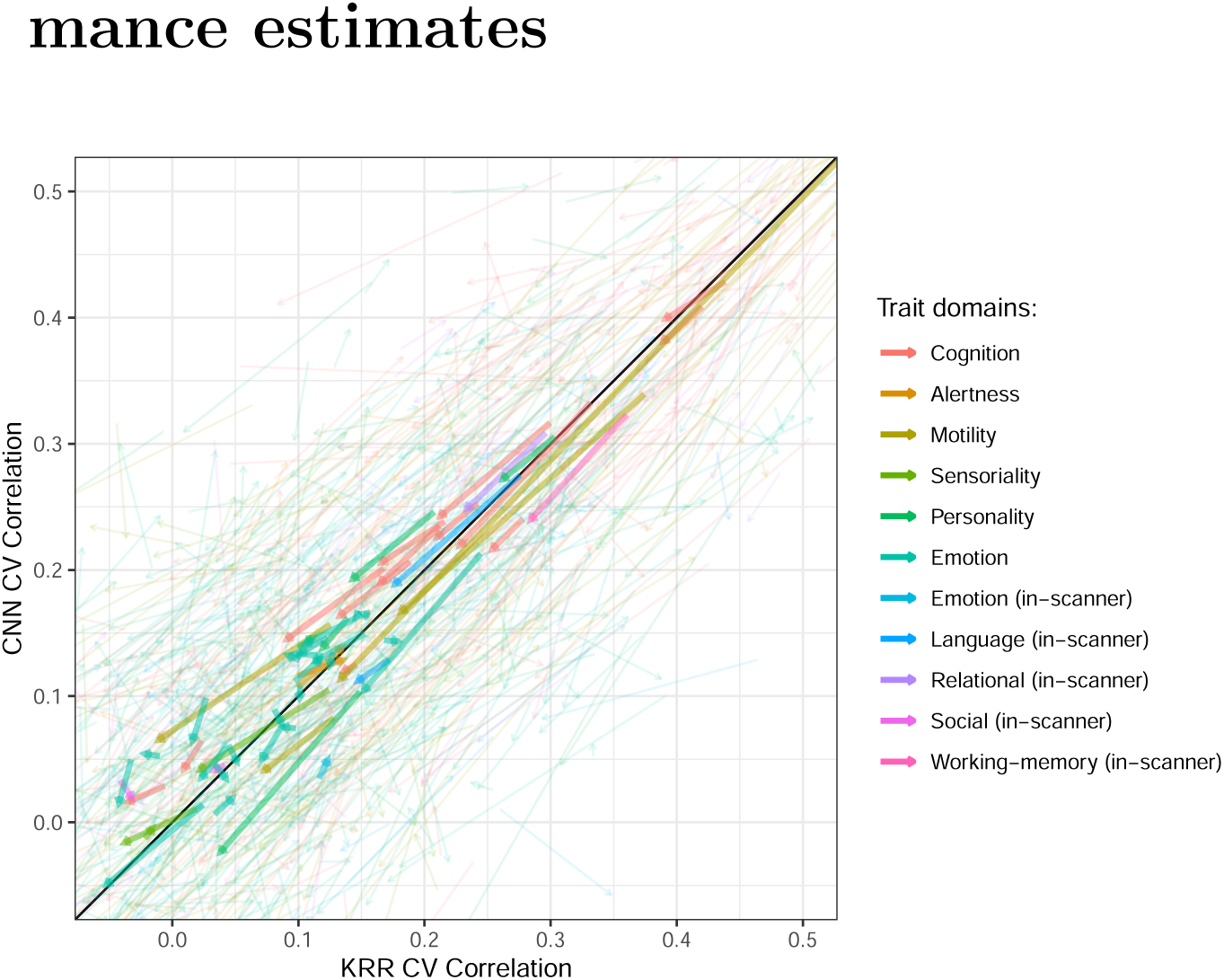
CV results in terms of Pearson’s correlation coefficient before and after deconfounding. Arrows start at the performance estimate without deconfounding and end at the performance estimate after deconfounding, per fold and trait. Thicker arrows denote averaged performance estimates across folds.

## G Task-performance measures and Self-reported measures

**Figure S12:**
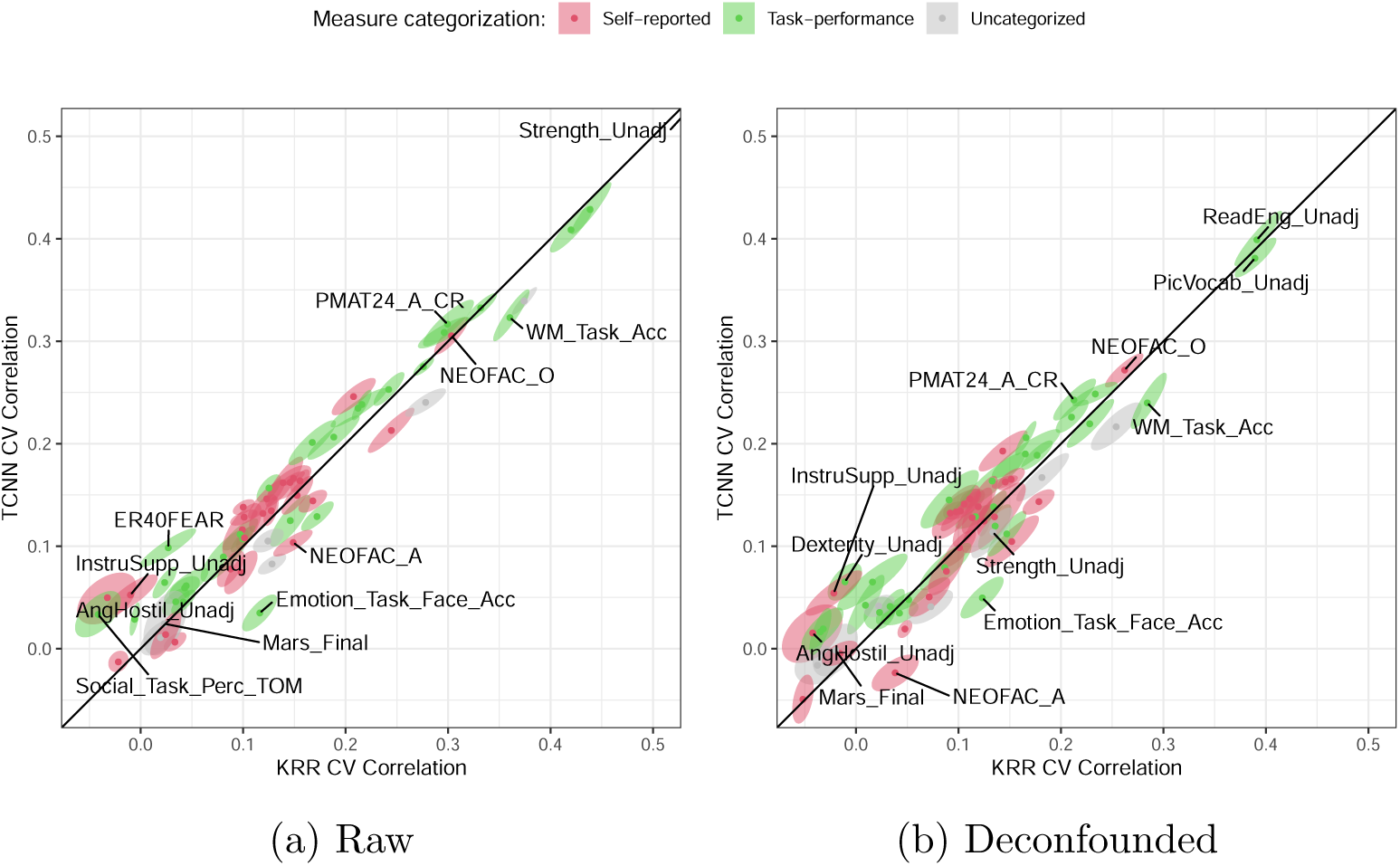
CV results in terms of Pearson’s correlation coefficient. (a) shows results without deconfounding while (b) includes deconfounding of predicted values and true labels. Ellipses delineate the 0.995 confidence region for the average based on bivariate t-statistics, not corrected for FCR. Behavioral scores colored green denote task-performance measures while behavioral scores red denote self-reported measures, according to Li et al. [6] and Líegeois et al. [43].

## Footnotes

1 The ten tests that compose the general factor in Dubois et al. [1] are: PicVocab – Unadj, ReadEng Unadj, CardSort Unadj, Flanker Unadj, ProcSpeed Unadj, VSPLOT – TC, PMAT24 A CR, PicSeq Unadj, ListSort Unadj, IWRD TOT.

